# Early Computational Detection of Potential High Risk SARS-CoV-2 Variants

**DOI:** 10.1101/2021.12.24.474095

**Authors:** Karim Beguir, Marcin J. Skwark, Yunguan Fu, Thomas Pierrot, Nicolas Lopez Carranza, Alexandre Laterre, Ibtissem Kadri, Abir Korched, Anna U. Lowegard, Bonny Gaby Lui, Bianca Sänger, Yunpeng Liu, Asaf Poran, Alexander Muik, Ugur Sahin

## Abstract

The ongoing COVID-19 pandemic is leading to the discovery of hundreds of novel SARS-CoV-2 variants on a daily basis. While most variants do not impact the course of the pandemic, some variants pose a significantly increased risk when the acquired mutations allow better evasion of antibody neutralisation in previously infected or vaccinated subjects or increased transmissibility. Early detection of such high risk variants (HRVs) is paramount for the proper management of the pandemic. However, experimental assays to determine immune evasion and transmissibility characteristics of new variants are resource-intensive and time-consuming, potentially leading to delays in appropriate responses by decision makers. Here we present a novel *in silico* approach combining spike (S) protein structure modelling and large protein transformer language models on S protein sequences to accurately rank SARS-CoV-2 variants for immune escape and fitness potential. These metrics can be combined into an automated Early Warning System (EWS) capable of evaluating new variants in minutes and risk-monitoring variant lineages in near real-time. The system accurately pinpoints the putatively dangerous variants by selecting on average less than 0.3% of the novel variants each week. With only the S protein nucleotide sequence as input, the EWS detects HRVs earlier and with better precision than baseline metrics such as the growth metric (which requires real-world observations) or random sampling. Notably, Omicron BA.1 was flagged by the EWS on the day its sequence was made available. Additionally, our immune escape and fitness metrics were experimentally validated using *in vitro* pseudovirus-based virus neutralisation test (pVNT) assays and binding assays. The EWS flagged as potentially dangerous all 16 variants (Alpha-Omicron BA.1/2/4/5) designated by the World Health Organisation (WHO) with an average lead time of more than one and a half months ahead of them being designated as such.

**One-Sentence Summary:** A COVID-19 Early Warning System combining structural modelling with machine learning to detect and monitor high risk SARS-CoV-2 variants, identifying all 16 WHO designated variants on average more than one and a half months in advance by selecting on average less than 0.3% of the weekly novel variants.

## Introduction

Despite a relatively slow mutation rate in human coronaviruses, since the emergence of the human coronavirus SARS-CoV-2 in Wuhan in December 2019, over 380,000 different missense variants (as of May 18, 2022) have been identified in the protein-coding viral sequences deposited in the GISAID ^1–3^ (Global Initiative on Sharing All Influenza Data) database and associated with multiple lineages. Of these, over 11,000 individual missense mutations (including indels) have been observed in the spike (S) protein, the key target for antibody neutralisation. While most mutations either reduce the overall fitness of the virus or bear no consequence to its features, some individual or combinations of mutations lead to high risk variants (HRVs) with modified immune evasion capabilities and/or improved transmissibility. For example, the Alpha (B.1.1.7, Q) variant of concern (VOC) spread widely due to higher transmissibility compared to the Wuhan variant, while the Beta (B.1.351) VOC has been shown to be less effectively neutralised by both convalescent sera and antibodies elicited by approved COVID-19 vaccines^4^. The Delta (B.1.617.2, AY) variant – characterised by high transmissibility – led to increased mortality and triggered a renewed growth in cases in countries with both high and low vaccination rates such as the United Kingdom^5^ and India^6^. More recently, the heavily mutated Omicron (BA.1) variant was amongst the quickest variants to be designated as a VOC by the WHO (World Health Organisation) due to a combination of widespread dissemination and several concerning mutations in the S protein as well as in other proteins^7^. Its descendants, BA.2 and BA.4/5, have acquired additional mutations in other neutralising antibody binding sites, allowing them to further escape host immune responses while simultaneously improving their ability to spread in the human population ^8–10^.

Hundreds of new variants are sequenced daily, only some of which are added to GISAID and other databases^2,11^. As new sequences continue to naturally emerge, the potential for the generation of variants that are both fit and highly immune resistant creates a significant challenge for public health authorities. The transmissibility and immune escape potential of a given variant could be assessed experimentally: evaluating one aspect of the fitness of variants requires experimental measurements of their binding affinity with its human receptor, angiotensin-converting enzyme 2 (ACE2), which is necessary for host cell infection; assessing immune escape potential requires *in vitro* neutralisation tests involving serum from vaccinated subjects or serum from patients previously infected with other variants of SARS-CoV-2. Both methods are resource-intensive and time-consuming and cannot be scaled to properly address the multitude of emergent variants.

In this work, we present a new method to evaluate SARS-CoV-2 variants based on *in silico* structural modelling and artificial intelligence (AI) language modelling and demonstrate that it captures features of a given variant’s fitness as well as its immune escape properties (Fig. 1.a). This approach is used to build an Early Warning System (EWS) that trains on the complete (up to a chosen time point) GISAID SARS-CoV-2 sequence database in less than a day and can score novel variants within minutes. It is a non-trivial task, as newly emerging HRVs most often comprise new sets of mutations and not all combinations of mutations present in previously identified concerning variants actually lead to enhanced immune evasion or transmissibility. The EWS is fully scalable as new variant data become available, allowing for the continuous risk monitoring of variant lineages. We show that it can flag HRVs weeks or even months earlier than their designation as such by the WHO, providing an opportunity to analyse subsequent mutations and variants in advance, thereby shortening the response time of health authorities ^12^.

**Fig. 1.**
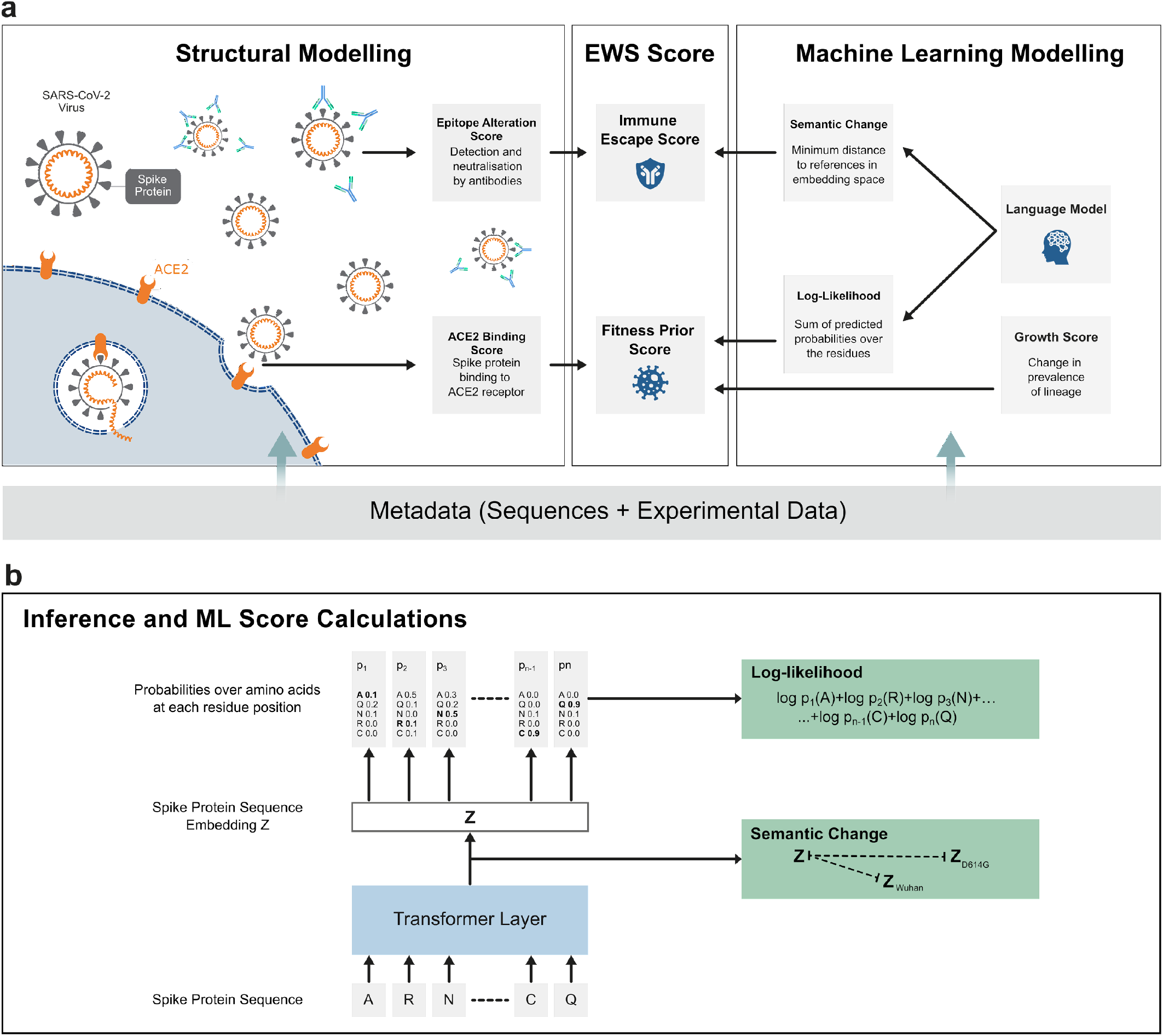
A schematic of the Early Warning System (EWS): structural modelling methods and natural language processing techniques to enable risk-level estimation of SARS-CoV-2 variants in real-time. **(a)** On the left hand side, structural modelling is used to predict the binding affinity of the SARS-CoV-2 S protein to host protein, ACE2, and to score the mutated epitope regarding its impact on immune escape. The right hand side panel represents the machine learning-based modelling that is used to extract implicit information from unlabeled data for the hundreds of thousands of registered variants in the GISAID database. Looking at the middle panel, the EWS relies on the information from structural modelling and machine learning modelling to compute an *immune escape score* and a *fitness prior score*, which taken together, present a more comprehensive view of the SARS-CoV-2 variant landscape. **(b)** Schematic of the machine learning model structure for assessing semantic change and log-likelihood. Once trained (Fig. S1a), the model receives as input a variant S protein sequence and returns an embedding vector of the S protein sequence as well as probabilities over amino acids for each residue position (Fig. S1b). The embedding vector is used to calculate the semantic change from reference variants such as Wuhan and D614G while the probabilities are used to compute the log-likelihood.

## Results

### *In silico* prediction of immune escape potential and *in vitro* validation

Mutations in the S protein are key to the heightened resistance to antibody-mediated neutralisation of new SARS-CoV-2 variants^13^. To evaluate the impact of said mutations on humoral immune evasion, 719 binding epitopes observed in 332 previously resolved structures of neutralising antibodies (nAbs)^14–17^ were mapped onto the S protein based on publicly available 3D structures (Table S1). An overlay of all nAb:S protein interaction interfaces was used to generate a colour-coded heatmap, indicating which surface-exposed amino acids are located in high epitope density regions (Fig. 2a). The number of unique nAbs whose binding epitope is affected by distinct SARS-CoV-2 variants’ mutations, normalised with respect to the total number of nAbs considered, was defined as the *epitope alteration score* (Table S2).

**Fig. 2.**
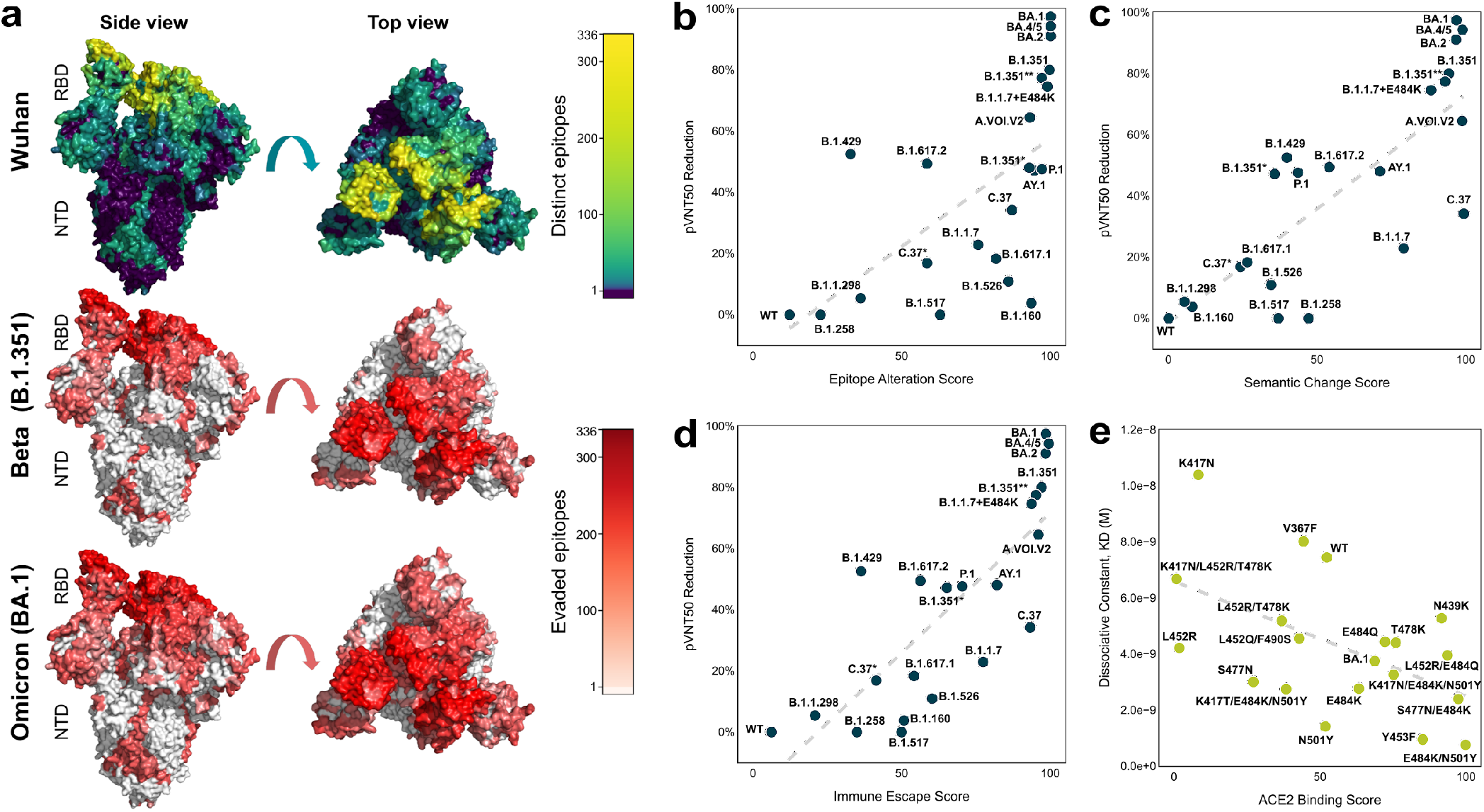
*In silico* predicted scores for immune escape and fitness prior correlate with *in vitro* data. **(a)** The surface of a SARS-CoV-2 S protein in the ‘one RBD up’ conformation (PDB ID: 7KDL) where the left-hand column shows a side view and the right-hand column shows a top-down view. The top row structure is coloured by the frequency of contact of surface residues with neutralising antibodies (brighter, warmer colour corresponds to more antibody binding). The middle and bottom rows depict the number of evaded epitopes in Beta (B.1.351) and Omicron (BA.1). **(b-d)** Validation of the immune escape metric with pseudovirus-based virus neutralisation test (pVNT) assay results. Relationships of the epitope alteration score, semantic change score, and combined immune escape score with the observed 50% pseudovirus neutralisation titer (pVNT_50_) reduction are shown across n=21 selected SARS-CoV-2 variants, including Omicron BA.1, BA.2, and BA.4/5, and the wild-type, Wuhan variant (WT). Cross-neutralisation of n=12-40 BNT162b2-immune sera was assessed against vesicular stomatitis virus (VSV)-SARS-CoV-2-S pseudoviruses. The pVNT_50_ reduction compared to wild-type SARS-CoV-2 (Wuhan variant) spike-pseudotyped VSV is given in percent. Variants for which experimentally measured geometric mean pVNT_50_ increased compared to the Wuhan variant were assigned a pVNT_50_ reduction of 0 (equal to wild type). Epitope alteration score (based on structural modelling) indicates the number of known neutralising antibodies with a known structure (max. n=227) whose binding epitope is affected by the SARS-CoV-2 variants’ mutations. Semantic change score (based on machine learning) indicates the predicted variation in the biological function between a variant and wild-type SARS-CoV-2, by calculating their L1 distance in embedding space. The scores were ranked and scaled in the range of [0, 100] such that the most immune escaping sequence has a score of 100. The immune escape score was calculated as the average of the scaled epitope alteration score and the scaled semantic change score. Dashed lines represent the fitted ordinary least squares regression line. **(e)** Validation of the ACE2 binding score with the experimentally determined ACE2 binding affinity (K_D_, dissociation constant) are shown across n=19 RBD variants, along with a fitted regression dashed line. The ACE2 binding score was ranked and scaled such that variants with the lowest and highest energy are assigned a score of 100 and 0, respectively.

Among all mutations, the ones that occur in the receptor-binding domain (RBD) are of particular importance due to this region’s direct interaction with the human ACE2 receptor, which both determines the propensity of a variant to infect a cell and its susceptibility to neutralising antibodies^18–20^. As seen in Table S3 and S4, the antibody epitopes used to calculate the epitope alteration score are well distributed across different RBD-targeting antibody classes defined by Barnes *et al.^21^* and epitope classes defined by Sikora *et al. ^22^*. However, the SARS-CoV-2 cell entry pathway and the neutralising antibody response against S protein is a complex process which involves other S domains as well, including the N-terminal domain (NTD) and cleavage sites ^23^. Thus, it is essential for the epitope alteration score to cover positions across the entire S protein^22^ (Table S5).

While the epitope alteration score can be used as a first proxy to evaluate escape from humoral immunity, it is limited by its dependence on the availability of antibody structure data. However, it was recently demonstrated that deep learning language models have the ability to capture the biological properties of proteins through unsupervised learning on large amounts of biological data^24–26^. At the inference step, the language model returns the predicted probability distribution of the 20 natural amino acids for each position in the protein (Fig. 1b), thereby leveraging the underlying biology of the large number of sequences seen during training from an evolutionary point of view. Hie *et al^27^* showed that language models trained on a dataset of proteins can be used to assess the risk of a viral variant. This risk was measured through the proxy known as *semantic change* (Table S2) to assess antigenic variation. In the work presented herein, the recurrent neural networks of Hie *et al^27^* were swapped for attention-based models (namely transformers)^28^ thereby replacing the auto-regressive approach to training the model used in Hie *et al^27^* with the Bidirectional Encoder Representations from Transformers (BERT) protocol. The BERT protocol has been shown to give better results in many applications^29^. In order to learn more general features of protein sequences and address unseen viral variants, the model was first pre-trained over the large collection of diverse proteins included in UniRef100 (non-redundant sequence clusters of UniProtKB and selected UniParc records) (Fig. S1a). The transformer model was then fine-tuned each month on the variants registered in GISAID (up to 125,000 unique S protein sequences vs. 4,172 S protein sequences in Hie *et al*.^27^), which is limited in scope to SARS-CoV-2. Additionally, our semantic change calculation differed from the one presented in Hie *et al^27^* in that it was computed to estimate the minimum change with respect to the reference sequences including the Wuhan variant, D614G variant and designated VOC (Fig. 1b).

In order to validate the immune escape *in silico* metrics, semantic change and epitope alteration score, *in vitro* pseudovirus-based virus neutralisation test (pVNT) assays were conducted (see Methods: VSV-SARS-CoV-2 S pseudovirus neutralisation assay). The cross-neutralising effect of n≥12 BNT162b2-immune sera was assessed against vesicular stomatitis virus (VSV)-SARS-CoV-2-S pseudoviruses bearing the S protein of 21 selected HRVs, including Omicron BA.1, BA.2, and BA.4/5 (Fig. 2b-d, Fig. S2, Table S6)^30–32^. The SARS-CoV-2 Omicron pseudoviruses were by far the most immune escaping compared to the other HRVs included in the experiment, with up to > 20-fold reduction of the 50% pseudovirus neutralisation titer (pVNT_50_) compared with the geometric mean titer (GMT) against the Wuhan reference spike–pseudotyped VSV ^32^. The calculated geometric mean ratio with a 95% confidence interval (CI) of the Omicron BA.1 pseudotype and the Wuhan pseudotype GMTs was 0.025 (95% CI; 0.017 to 0.037), indicating another 10-fold drop in the neutralising activity against Omicron BA.1 compared to the second most immune escaping B.1.1.7+E484K pseudovirus with a geometric mean ratio of 0.253 (95% CI; 0.196 to 0.328) (Fig. S2c). This result is in good concordance with the *in silico* immune escape score for Omicron BA.1 which was the highest amongst observed, circulating variants at the time of performing the assay. Since then, new Omicron subvariants have surpassed the ancestral BA.1/BA.2 lineages in terms of EWS metrics.

Across all HRV pseudoviruses tested, the epitope alteration score correlates positively with the calculated pVNT_50_ reduction (Fig. 2b; Pearson *r*=0.64, *p*=1E-3; Spearman *r*=0.77, *p*=3E-5). In particular, we observed that the predictive power of the epitope alteration score is distributed across different antibody and epitope classes rather than being dependent on a single class (Table S7-S9). Positive correlation with pVNT_50_ reduction was also observed with the semantic change score (Fig. 2c; Pearson *r*=0.81, *p*=5E-6; Spearman *r*=0.76, *p*=3E-5). Furthermore, an average of both *in silico* scores (summarised as the *immune escape score*) exhibits an even stronger correlation with the observed reduction in neutralising titers (Fig. 2d; Pearson *r*=0.81, *p*=5E-6; Spearman *r*=0.86 *p*=3E-7). This *immune escape score* was subsequently further compared with multiple independent studies^4,33–37^, comprising data from individuals vaccinated with different, approved COVID-19 vaccines. Consistent strong correlations have been observed with respect to the independently performed experimental assays thereby demonstrating the accuracy and robustness of this score (Fig. S3-S5).

### *In silico* estimation of fitness and *in vitro* validation

The immune escape score predicts if a given viral variant may evade neutralisation by the immune system, but it does not capture protein changes that either enhance the efficacy of viral cell entry or negatively impact its structure or function. Capturing the full transmissibility potential of the virus (*fitness*) is beyond the scope of this work as it comprises many complex dynamics, however, we can propose three informative priors contributing toward it: *ACE2 binding score, conditional log-likelihood score* and *growth score* (Table S2).

A key determinant of viral spread is the effectiveness with which virus particles can attach to and invade host cells. This characteristic is especially important when considering individuals without pre-existing immunity or viral variants which are able to better evade immune responses. In order to infect the human host cell, the RBD of the viral S protein associates with ACE2, the cellular receptor for SARS-CoV-2. Therefore, we assessed the fitness prior based on the predicted impact of sets of mutations on the binding affinity of the variant S protein to the human ACE2 receptor, herein referred to as the *ACE2 binding score*. The interaction between a variant S protein and the ACE2 protein was computed through repeated, fully flexible *in silico* docking experiments (see Methods: ACE2 binding score), allowing for an unbiased sampling of the binding landscape. In order to reduce the required computational resources, the S protein modelling was restricted to its RBD domain, i.e., the domain known to be directly binding to the ACE2 receptor. To calculate the ACE2 binding score, we used the median binding energy (the change in estimated Gibbs free energy, ΔG, between bound and unbound states), which acts as a proxy for global complex affinity (Fig. S6).

In order to assess the validity of the ACE2 binding score, the simulation results were compared with *in vitro* results. Surface plasmon resonance (SPR) kinetic analysis was performed to determine the affinity (K_D_, dissociation constant) of 19 RBD variants to the ACE2 receptor (see Methods: Binding kinetics of RBD variants to ACE2 using surface plasmon resonance spectroscopy). Notably, this assay measures observable association rates, which are a result of a dynamic process, whereas simulations measure aggregated, static binding affinity, thus marginalising the contribution of mutations toward the flexibility and kinetics of the S protein. Despite this, the ACE2 binding score used herein shows meaningful correlation with the K_D_ values (Fig. 2e; Pearson *r*=-0.53, *p*=2E-2; Spearman *r*=-0.45, *p*=5E-2).

Another aspect that partially models the fitness of a variant is how similar a given variant is to other variants which have been known to exhibit rapid population growth. This analysis is not achievable through simple sequence comparison due to epistatic interactions between sites of polymorphism in which certain mutation combinations enhance fitness while being deleterious when they occur separately. Therefore, the same trained transformer model described previously was leveraged to calculate the log-likelihood of the input sequence. From a language model perspective, the higher the log-likelihood of a variant, the more probable the variant is to occur. In particular, the log-likelihood metric supports substitutions, insertions, and deletions without requiring a reference sequence to measure against, unlike the *grammaticality* of Hie *et al*.^27^ which requires a reference sequence. Our language model was not provided with explicit sequence count data in the training phase, yet on average our language model assigned higher log-likelihood values to sequences with a higher actual observed count (Fig. S7; Pearson *r*=0.96, *p*=1E-11; Spearman *r*=0.99, *p*=3E-20). High log-likelihood may indicate features common in the general variant population, which are likely to be fitness-related, thus allowing variants harbouring these to sustain additional such mutations.

However, the values of log-likelihood tend to diminish with the increasing number of mutations, which is expected given the definition of this metric; this creates a bias toward variants with low mutation counts. Considering that all the samples used for training have been detected in patients, and as such have likely satisfied minimal fitness criteria, we addressed this bias by introducing the *conditional log-likelihood score*, measuring how the log-likelihood of the variant in question compares to other variants with similar mutational loads as opposed to the entire viral population. This metric sheds more light on highly mutated, potentially concerning variants like Omicron BA.1. Due to its high mutational load, Omicron BA.1 may be perceived by raw log-likelihood as highly unlikely, however, relative to other variant sequences with a similar number of mutations, it becomes clear that Omicron BA.1 stands out, leading to a higher conditional log-likelihood score (Fig. S8).

None of the metrics discussed above capture the entirety of factors affecting the frequency of viral variants. Additionally, conditional log-likelihood is a metric measuring similarity to already known, rapidly growing variants. By its nature, it cannot fully assess variants that exhibit completely new sequence features (until these features are observed more often). Therefore, the fitness prior metric includes *growth score*, an empirical term representing the quantified change in the fraction of the sequences in the database that a variant in question comprises. This addresses the intuitive notion that variants which are increasing in prevalence are more imminently interesting than those which are not. The growth metric also reflects the overall fitness of a given variant, implicitly taking into account the impact of mutations beyond the RBD domain and S protein. Furthermore, it complements the ACE2 binding score which models only the RBD domain.

### Combining fitness prior and immune escape scores to continuously monitor high risk variants

Different selective pressures on virus evolution lead to variants with high immune escape and fitness because a virus must remain evolutionarily competent to successfully spread. We hypothesise that a system that keeps track of immune escape and fitness factors (as depicted in Fig. 3 and Fig. S9) could continuously monitor HRVs on a near real-time basis as new sequences are evaluated and added to the data pool in a matter of minutes. To begin probing this hypothesis, for each week corresponding to the VOC designation, we averaged the immune escape score and fitness prior score per lineage in order to rank and visualise the changing landscape of the variants. It is important to note that the ranking of any sequence – and consequently lineage – is relative and depends on the scores of the other circulating variants. While relative ranking of individual variants with regard to each metric remains the same, the ranking values differ from week-to-week. Therefore, compound scores (like immune escape) tend to differ as well which allows for the granularity necessary to discriminate between HRVs in different circumstances, for example, when there are multiple variants circulating, each with its own characteristics, or when the landscape is dominated by subvariants of a single lineage (e.g., the Delta outbreak of Summer 2021). As seen in Fig. 3, the VOCs are mostly located relatively high in the upper-right corner at the week of WHO designation, i.e., they are comparatively highly immune escaping and have satisfactory fitness prior score for their immune escape value. Through time, the prevalent VOCs, such as Alpha: B.1.1.7, Delta: B.1.617.2 and AY, Omicron: BA.1 and BA.2, often diversify and lead to the emergence of sub-lineages. These sub-lineages often have increased fitness prior at the cost of decreased immune escape score for improved transmissibility. Simultaneously, with new emerging VOCs becoming more prevalent, the past VOCs prevalence gradually decreases with reduced scores for both fitness prior and immune escape.

**Fig. 3.**
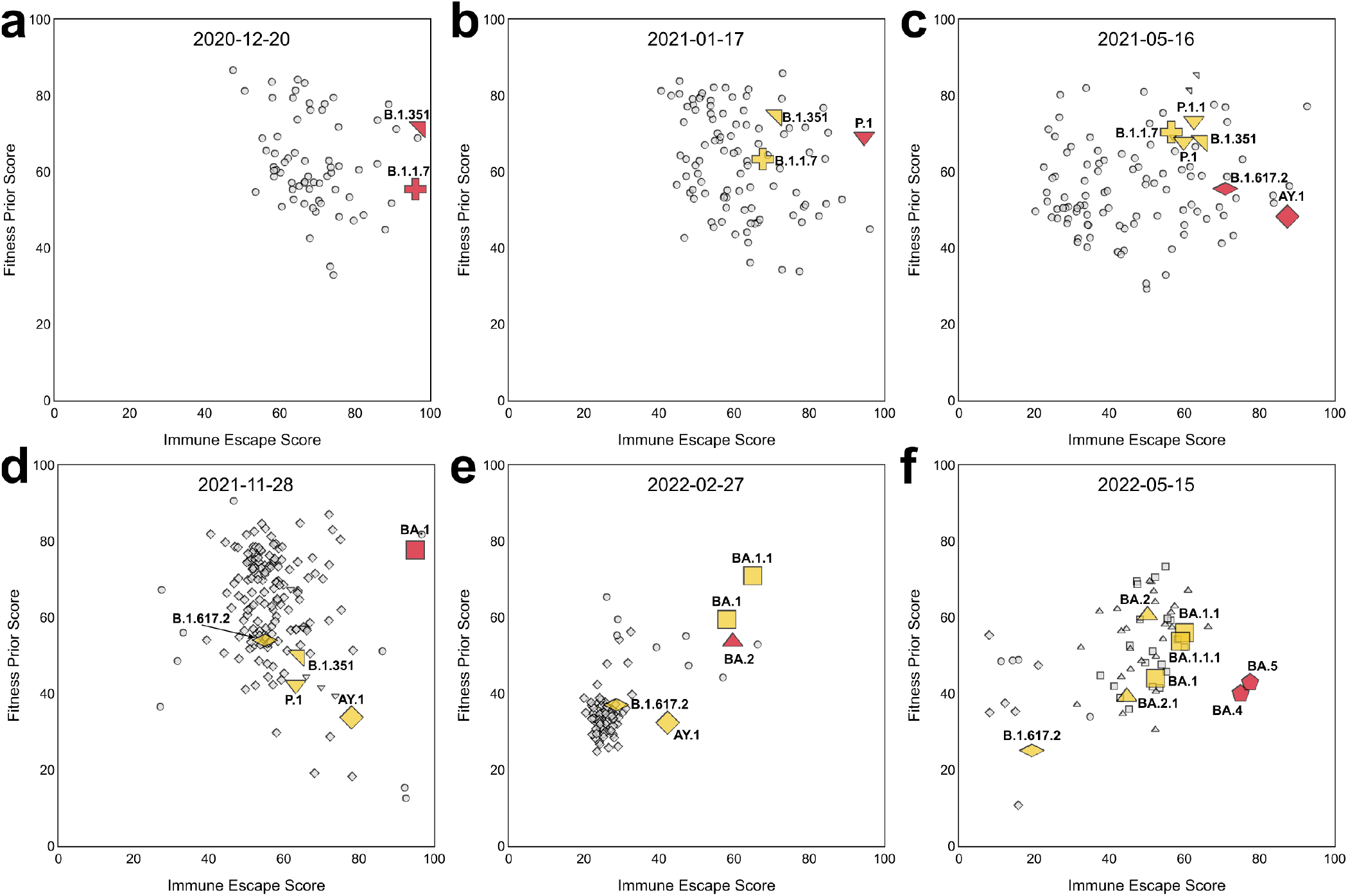
Combining immune escape and fitness prior for continuous monitoring. Snapshot of lineages in terms of fitness prior and immune escape score on **(a)** December 20th 2020, **(b)** January 17th 2021, **(c)** May 16th 2021, **(d)** November 28th 2021, **(e)** February 27th 2022, and **(f)** May 15th 2022, corresponding to the week of the designations of Alpha/Beta, Gamma, Delta, Omicron BA.1, Omicron BA.2 and Omicron BA.4/BA.5. Red markers indicate the designated lineages of the week, yellow markers are the previously designated lineages and grey markers indicate other lineages. Circles correspond to non-VOC lineages and other symbols correspond to designated variants and their closely related lineages. The cross, north-east-pointing triangle, south-pointing triangle, flattened diamond, diamond, square, north-pointing triangle, and pentagon correspond to Alpha (B.1.1.7), Beta (B.1.351), Gamma (P.1), Delta (B.1.617.2, AY), Omicron BA.1, BA.2, and BA.4/BA.5 lineages, respectively. Lineages with the same S protein sequence are indicated with the same shape (e.g., BA.4 and BA.5 are pentagons). Only lineages that have been observed within the past 8 weeks relative to the indicated date for each plot and have been reported more than 10 times are included. Note how initially innocuous BA.2 lineage(e), has evolved into highly immune evasive BA.5 subvariant (f). See Fig. S9 for the corresponding density contour plots of sequences.

### Detection of potentially high risk variants prior to substantial spread in the population

Experimental assays aiming to determine a given variant’s immune evasion and fitness are time and resource-intensive. Available data show that thousands of new variants are emerging each week at an increasing rate (on average 70 per week in September 2020, 6,500 in August 2021 and 11,500 in February 2022 in GISAID database). Moreover, this number is likely an underestimate given limited viral sequencing and data deposition in many countries. It is not feasible for health authorities to perform preventative experimental assessments whenever a new variant is identified, despite the benefits of a proactive stance in detecting HRVs before their spread.

As shown in the previous section, the VOCs are often among the top immune escaping lineages, while the fitness prior scores are not necessarily the highest as observed with, for example, Delta (B.1.617.2, AY) in May 2021 (Fig. 3c). Furthermore, it was recently proposed that prolonged viral evolution in immunocompromised patients who are not able to clear the virus generates intrapatient viral variants with increased immune evasion rather than increased fitness ^38–41^. Some of the new variants reside on potentially long branches of the viral phylogenetic tree ^42^, indicating the lack of information about the intermediate ancestors of these variants. These results, together with increased vaccination rates worldwide, put an added emphasis on immune evasion as a key risk factor in newly emerging variants, which motivated our approach to only use immune escape for early HRV detection.

To address this issue, we present an *in silico* EWS built to scan sequences at scale which would allow for the selection of a limited number of only the most predicted high-risk sequences for experimental exploration and validation each week^12^. On a weekly basis, the EWS scores and ranks all of the sequences reported in the past eight weeks. The non-VOC, novel sequences of the week are then selected for further analysis. To increase the sequence diversity while reducing the sequence pool size, the sequences are clustered by their RBD mutations and only the top immune escaping sequence is kept per RBD variant. Retaining additional sequences per RBD variant does not impact the utility of the system, but may potentially lead to selecting multiple, largely equivalent variants. Lastly, the top 12 immune escaping sequences are selected. The selected sequences are subsequently aligned and used to populate a heatmap to highlight potential concerning mutations and sequences each week to be validated with experimental studies. The overview of this detection protocol is presented in Fig. 4.

**Fig. 4.**
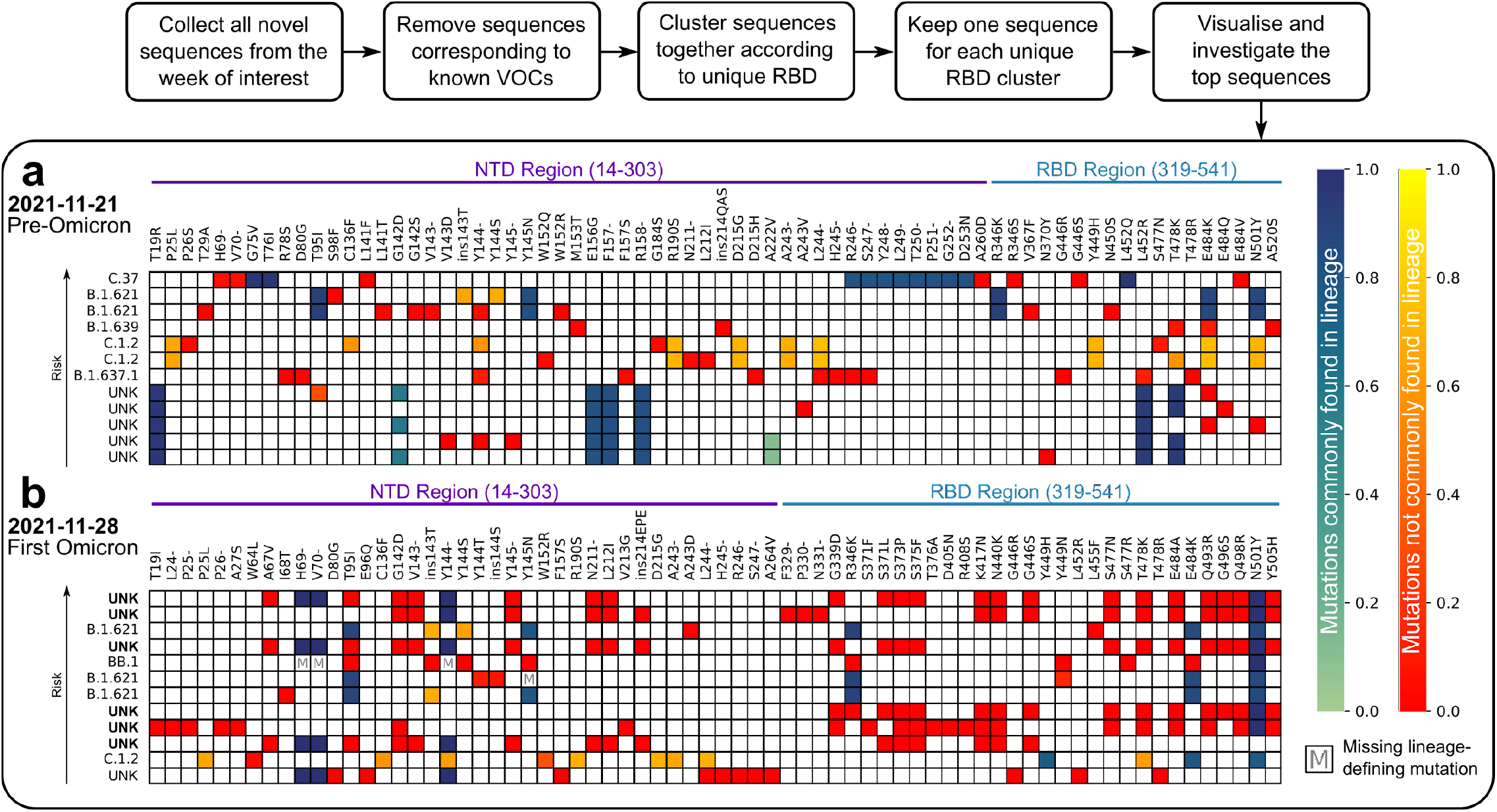
EWS detection protocol and weekly report. A heatmap is constructed each week with the top selected sequences for the immune escape score resulting from the detection protocol outlined above. The two heatmaps shown here represent the top sequences resulting from this protocol from the week of November 21, 2021 **(a)** and November 28, 2021 **(b)**, the weeks corresponding to the time just prior to and the week of the appearance of Omicron in the global population, respectively. Each row represents a sequence in decreasing order of immune escape score from top (highest score) to bottom (lower score). The label at the front of each row indicates either an associated, reported lineage or – as indicated by the UNK label – an unknown lineage. The UNK labels in bold in (b) indicate sequences that were later designated as belonging to Omicron. Sequences labelled with UNK were compared to the closest VOC lineage based on sequence similarity in order to distinguish common and uncommon mutations as indicated in the colourbars. All non-bolded UNK-labelled sequences in (a) and (b) were later designated to other lineages. Each column represents a mutation present in the S protein within the NTD and RBD regions indicated in purple and teal, respectively. The most significant colours overall are green and red. The colour scale from green to blue represents lineage-defining mutations (defined by WHO or inferred from mutation frequency) and the relative frequency of a mutation from 0.0-1.0 in the sequence population of the lineage. Green is indicative of mutations that are less essential than those indicated in blue. The colour scale from red to yellow represents non-lineage-defining mutations and their frequency from 0.0-1.0 in the sequence population of the lineage. Red mutations have been seen less frequently (more novel). Boxes marked with an M indicate lineage-defining mutations that are missing from the corresponding sequence.

We noted a striking difference between the heatmap corresponding to the week of Omicron BA.1 lineage emergence and that immediately preceding it. The pre-Omicron week (Fig. 4a) is characterised by incremental changes to existing lineages in which already circulating, fitness-inducing mutations (like deletions in NTD loops) are observed. The low number of unique mutations and their relatively high background frequency (*heat* on the red-yellow scale), are diverse across samples, and thus are not imminently threatening. The samples with unassignable lineages (UNK) are ranked low and appear to be slight variants of circulating lineages harbouring inconsequential mutations. In the week of Omicron BA.1 emergence (Fig. 4b), we observed a large fraction of new, previously unobserved mutations (in deep red). Most of the unassignable lineage samples, which were later assigned as Omicron BA.1 variants (in bold in Fig. 4b), are ranked high and harbour the same clusters of mutations. These traits combined strongly suggest emergence of a new (sub)lineage of potential high risk.

### EWS flagged all 16 WHO designated variants months ahead

To assess the system’s precision, a retrospective analysis was conducted for every week between September 16th, 2020 and May 15th, 2022 using the detection protocol described in the previous section. For each week, the language model was trained on variants up to that week and any other data used were from prior to the analysis date. We focused on the detection of the 16 variants designated by the WHO (Alpha, Beta, Gamma, Delta, Epsilon, Zeta, Eta, Theta, Iota, Kappa, Lambda, Mu, Omicron BA.1, BA.2, BA.4 and BA.5). In particular, we were interested in detecting designated lineages before their WHO designation, which often represents a small proportion of the sequence candidates, as shown in Fig. 5a.

**Fig. 5.**
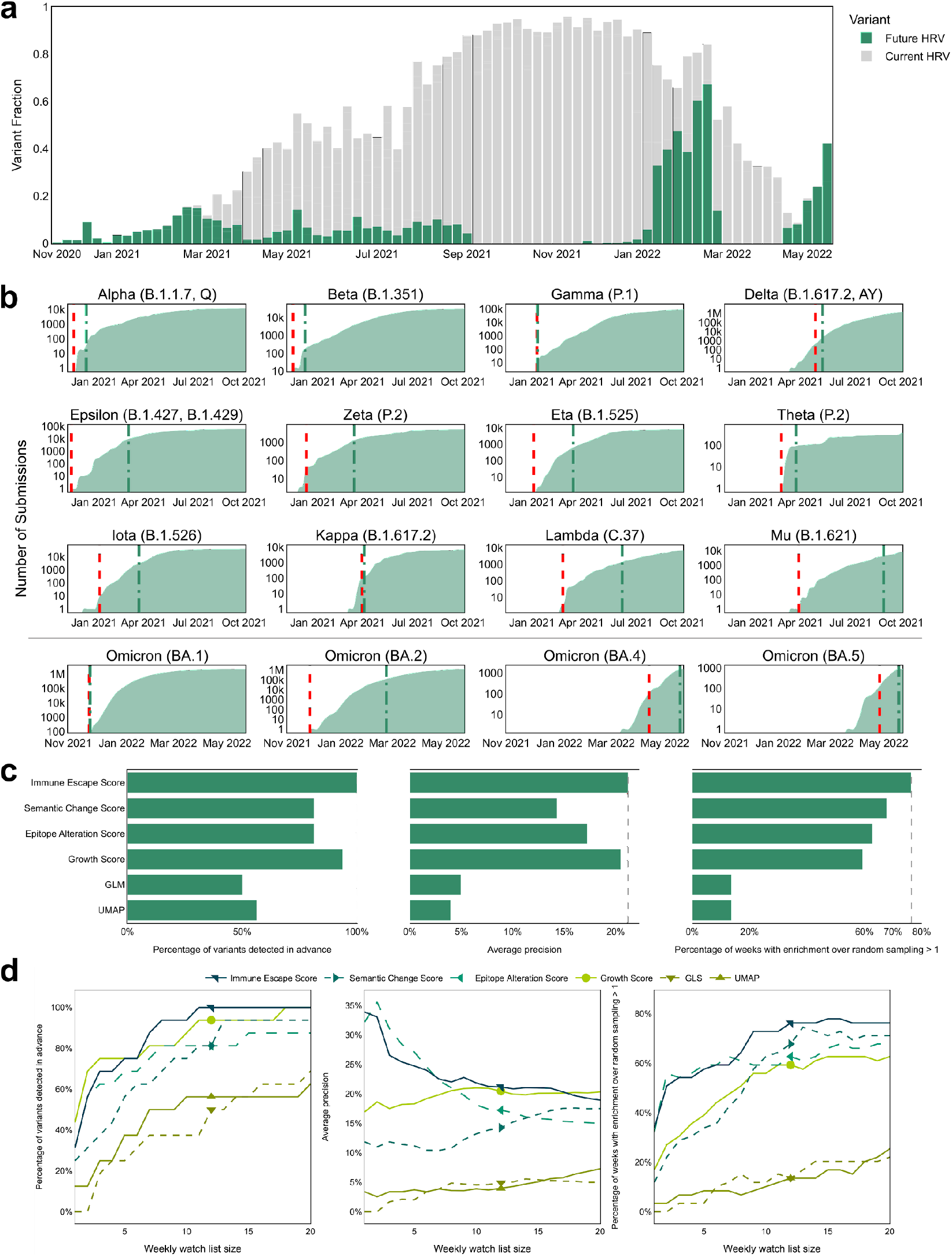
The EWS flags High Risk Variants ahead of their WHO designation. **(a)** The goal of the EWS is to detect the sequences each week that have not yet been designated as HRVs but will be designated as HRVs in the future. Each bar corresponds to a week and represents all novel, non-VOC sequences and it can be split into 1) sequences that will later be designated as HRVs, corresponding to the green portion 2) sequences that were labelled to belong to unknown lineages that later will be designated as known VOCs and, therefore, are still included in our novel, non-VOC set of sequences, corresponding to the grey portion, and 3) other sequences, corresponding to the white portion. **(b)** The cumulative sum of all cases of a given variant lineage (in log scale) over time. Vertical lines indicate the date of WHO designation of a given variant (green dot-dashed) vs. date of flagging by the EWS (red dashed) using a weekly watch-list size of 12 variants. **(c)** Detection results using immune escape score, semantic change score, epitope alteration score, growth score, generalised linear model (GLM) and UMAP model. The left bar chart displays the number of HRVs detected ahead of their designation. The centre bar chart displays the average precisions over the weeks. The right bar chart displays the proportion of the weeks where the used metric achieves enrichment (ratio between the number of correctly tagged HRV samples versus the number expected in random sampling) larger than one (better than random). **(d)** Detection results with respect to the watch list size per week using immune escape score, semantic change score, epitope alteration score, growth score, generalised linear model (GLM) and UMAP model. The markers correspond to 12 sequences per week. The immune escape score outperforms other methods consistently across all metrics.

When using a weekly watch list with a size of 12 variants (compared to the thousands of novel variants reported per week, the watch list corresponds to less than 0.3% of the data on average), the EWS flagged all 16 WHO designated variants (Fig. 5b) with an average lead time of 52 days (more than a month and a half) before these were designated as such by the WHO (Table S10). Specifically, the EWS identified Omicron BA.1 as the highest immune escaping variant over more than 70,000 variants discovered between early October and late November 2021. The particular combination of frequently occurring RBD mutations – with some unique to Omicron sub-lineages – and NTD indels, allows the virus to evade a large fraction of neutralising antibodies, which was accurately captured by the exceptionally high epitope alteration score (Fig. 3d-e). Roughly at the time of emergence of BA.1, another variant, BA.2, emerged (Fig. 3e). While early BA.2 variants were not as immune escaping as BA.1, several of its descendants, including BA.4 and BA.5 variants, were flagged by the EWS as increasingly immune escaping upon emergence, outperforming BA.1/BA.2 variants (Fig. 3f).

One can consider the growth score alone as a plausible metric that requires neither machine learning nor simulation to detect 15 HRVs early (Fig. 5c, Table S11). However, the immune escape score implemented in the EWS can detect all 16 HRVs and the growth score requires 50% more sequences per week to detect Gamma. Immune escape score also outperforms the growth score across the variants in terms of average lead time ahead of WHO designation (Table S11). Importantly, the immune escape score only requires the nucleotide sequence of the S protein, while the growth score requires additional observational data, such as occurrence frequencies. This makes the growth score applicable only in the case of already ongoing outbreaks or variant spread which precludes their application for newly emerging and theoretical variants, hence reducing its utility in real life situations (see Methods: Growth score). Moreover, we evaluated the system in terms of precision, which is equivalent to the true positive rate (1 - false positive rate) in the weekly watch list, where a positive class represents sequences that would later be designated as HRVs. Maximising precision requires the system to recall as many HRV sequences as possible with a fixed watchlist size; the growth score underperforms the immune escape score when only a limited number of sequences can be detected per week (Fig. 5d). In particular, the growth score fails to detect HRVs during the majority of the Summer of 2021 when many lineages were spreading widely (Fig. S10).

However, as the analysis used a fixed-length list of predictions each week, the typical classification metrics such as precision are limited as there are periods when there is nothing new to detect (thus limiting the precision). We compared the number of detected HRV sequences with the number expected from random sampling each week. The ratio is defined as an enrichment score, where 1 means the method detects as many as expected via random sampling, and larger than 1 means the method detects more HRVs than random sampling (see Methods: Enrichment score). We suggest that enrichment over the random baseline forms a much more informative metric in this context. The analysis demonstrated that the immune escape score consistently has more weeks with enrichment higher than 1 (better than random) than any other metric, including the growth metric (Fig. 5d). With detecting 12 sequences per week, across the 59 weeks where there exist HRVs to be designated, the immune escape score outperforms random sampling (achieves an enrichment score greater than 1) in 45 weeks (more than 75% of the time) while the growth score has only 35 weeks (less than 60% of the time) with a significant enrichment greater than 1 (*p*=2E-2; paired *t*-test).

The immune escape score used by the EWS to detect HRVs early combines the epitope alteration score and semantic change score (Fig. 1a). We evaluated the early detection performance of each one of these two components both individually and combined. The epitope alteration score and semantic change score both detect 13 out of 16 WHO designated variants ahead of time. Their combination, however, flags all WHO designated variants (Fig. 5c). We also applied standard machine learning techniques, both supervised and unsupervised (denoted “GLM” and “UMAP”, respectively) corresponding conceptually to epitope alteration score and semantic change score, respectively (see Methods: Detecting HRVs by standard ML methods). These standard machine learning techniques do not reach the same predictive performance as the methods proposed in this work (Table S11). This validates our approach of associating protein structure modelling and transformer language models on protein sequence to accurately rank SARS-CoV-2 variants.

## Discussion

Through validation of our immune escape and fitness prior scores using published and newly generated data, we show that the combination of structural simulations, machine learning and genomic sequencing of SARS-CoV-2 variants allows for large-scale continuous risk monitoring and precise early detection of HRVs.

Every attempt to predict real life events is fraught with predictive error. We have demonstrated that EWS is capable of detecting pertinent, ominous variants when they occur and, critically, prior to their WHO designation as such. Filtering down to a hit list of only 12 sequences per week from thousands (i.e., on average less than 0.3% of weekly novel variants), the Early Warning System can accurately detect all 16 HRVs months ahead of the official WHO designation. Additionally, detection is often possible within the same week the sequenced variant enters the database. For instance, the EWS flagged Omicron BA.1 as soon as it was uploaded to GISAID – based on its sequence alone – ranking it to be one of the highest immune escaping variants ever documented for SARS-CoV-2 variants. Compared with other baselines, including growth metrics, heuristics, and machine learning models, the EWS achieves significantly higher predictive power in terms of enrichment over random sampling and all other considered metrics, demonstrating a sustained precision and robustness over the period from 2021 to 2022.

Combining comprehensive sequencing with structural modelling and machine learning can provide unprecedented insights into the COVID-19 pandemic through detailed weekly reports and analysis of predicted top variants. Together with the experimental assays, these collections of sequences and mutations could be harnessed by public health authorities and governments worldwide to increase their preparedness for HRVs and potentially alleviate associated human and economic costs. Future development of our approach could include expanding the immune escape scores to consider varied vaccination status and developing further functionalities such as the assessment of known and predicted T-cell epitopes and the projection of prospective variant evolution scenarios.

## Methods

### Variant notations

We refer to a variant as a protein sequence of a coronavirus’ S protein that differs from the original Wuhan spike protein that we refer to as wild type. We represent a variant in terms of its mutations with respect to the Wuhan strain. For instance, the notation N501Y represents a substitution in position 501, replacing N with Y; the notation ins214EPE represents inserting EPE after position 214; and notation H69-V70-represents a deletion of H and V at positions 69 and 70, respectively.

### VSV-SARS-CoV-2 S pseudovirus neutralisation assay

A recombinant replication-deficient VSV vector that encodes a green fluorescent protein (GFP) and luciferase (Luc) instead of the VSV-glycoprotein (VSV-G) was pseudotyped with SARS-CoV-2 spike (S) protein derived from either the Wuhan reference strain (NCBI Ref: 43740568) or variants of interest according to published pseudotyping protocols^43^. The mutations found in S of the VOCs are listed in Table S6. In brief, HEK293T/17 monolayers transfected to express SARS-CoV-2 S with the C-terminal cytoplasmic 19 amino acids truncated (SARS-CoV-2-S[CΔ19]) were inoculated with the VSVΔG-GFP/Luc vector. After incubation for 1 hour at 37 °C, the inoculum was removed, and cells were washed with PBS before a medium supplemented with anti-VSV-G antibody (clone 8G5F11, Kerafast) was added to neutralise the residual input virus. VSV-SARS-CoV-2 pseudovirus-containing medium was collected 20 hours after inoculation, 0.2 μm filtered and stored at −80 °C.

For pseudovirus neutralisation assays, 40,000 Vero 76 cells were seeded per 96-well. Sera were serially diluted 1:2 in culture medium starting with a 1:15 dilution (dilution range of 1:15 to 1:7,680). VSV-SARS-CoV-2-S pseudoparticles were diluted in a culture medium to obtain either ~1,000 or ~200 transducing units (TU) in the assay. Same input virus amounts for all pseudoviruses were used within an individual experiment. In total 8 experiments were performed covering the SARS-CoV-2 variants listed in Table S6 always taking Wuhan S pseudovirus as reference. Serum dilutions were mixed 1:1 with pseudovirus for 30 minutes at room temperature prior to addition to Vero 76 cell monolayers and incubation at 37 °C for 24 hours. Supernatants were removed, and the cells were lysed with luciferase reagent (Promega). Luminescence was recorded, and neutralisation titres were calculated by generating a four-parameter logistic fit of the percent neutralisation at each serial serum dilution. The pVNT_50_ is reported as the interpolated reciprocal of the dilution yielding a 50% reduction in luminescence. If no neutralisation yielding a 50% reduction in luminescence was observed, an arbitrary titer value of 7.5, half of the limit of detection (LOD), was reported.

### Binding kinetics of RBD variants to ACE2 using surface plasmon resonance spectroscopy

Binding kinetics of RBD variants was determined using a Biacore T200 device (Cytiva) with HBS-EP+ running buffer (BR100669, Cytiva) at 25 °C. Carboxyl groups on the CM5 sensor chip matrix were activated with a mixture of 1-ethyl-3-(3-dimethylaminopropyl) carbodiimide hydrochloride (EDC) and N-hydroxysuccinimide (NHS) to form active esters for the reaction with amine groups. anti-mouse-Fc-antibody (BR100838, Cytiva) was diluted in 10 mM sodium acetate buffer pH 5 (30 μg/mL) for covalent coupling to immobilisation level of ~6,000 response units (RU). Free N-hydroxysuccinimide esters on the sensor surface were deactivated with ethanolamine-HCl.

Human ACE2-mFc (10108-H05H, Sino Biological Inc.) was diluted to 5 μg/mL with HBS-EP+ buffer and applied at 10 μL/min for 15 seconds to the active flow cell for capture by immobilised antibody, while the reference flow cell was treated with buffer. Binding analysis of captured hACE2-mFc to RBD variants (40592-V08B, 40592-V08H113, 40592-V08H115, 40592-V08H82, 40592-V08H59, 40592-V08H84, 40592-V08H85, 40592-V08H112, 40592-V08H28, 40592-V08H81, 40592-V08H90, 40592-V08H91, 40592-V08H88, 40592-V08H86, 40592-V08H111, 40592-V08H80, 40592-V08H1, 40592-V08H14, 40592-V08H46, 40592-V08H121 40592-V08H121, Sino Biological Inc.) was performed using a multi-cycle kinetic method with concentrations ranging from 3.125 to 50 nM. An association period of 120 seconds was followed by a dissociation period of 300 seconds with a constant flow rate of 30 μL/min and a final regeneration step. Binding kinetics were calculated using a global kinetic fit model (1:1 Langmuir, Biacore T200 Evaluation Software Version 3.1, Cytiva).

### Data

The S protein sequences were collected from GISAID up until May 18th, 2022. For each S protein sequence, the missing amino acids were filled using the next known amino acid and the lineage assignment was performed using PANGOLIN^44^. Mutations with respect to the wild type were calculated using Clustal Omega^44,45^ and HH-suite^24^.

Multiple data cleaning procedures were performed, including removing sequences not conforming to basic biological assumptions (e.g., lacking the initiating methionine), sequences having more than ten continuous mutations (indicating a very likely frameshift), and sequences whose submissions date is more than two months later than its collection date (as these may be valid, but often are incorrectly dated, thus affecting the benchmark accuracy of the system).

### Language modelling

The domain of Natural Language Processing (NLP) has experienced several breakthroughs in the past years. The emergence of recurrent and attention-based deep neural networks has led to impressive results for text generation and translation. Recently, this technology has been leveraged to learn the language of biology^25,26^. How it works can be described with a simple analogy where protein sequences are considered as sentences and amino acids as words. The models are trained on large datasets of known protein sequences in an unsupervised manner. In other words, there is no need to label the data and any newly registered protein sequence can be exploited.

Information about protein properties is stored at two positions inside the model once it is trained. On one side, the probabilities returned by the model indicate how likely this sequence is to be natural/viable/feasible. On the other hand, the outputs of the model’s layers – and notably the last layer – provide a high-dimensional representation for each sequence, which we call the embedding of the protein. The embedding of the protein contains information about the protein properties and can be used either directly or to train a classification or regression model. Recently, Meier *et al*. demonstrated that these models also capture the effects of mutations on protein function^46^.

### Model architecture

In this work, the input of the model consisted of the sequence characters corresponding to the amino acids forming the protein. Each amino acid was first tokenised, i.e., mapped to their index in the vocabulary containing the 20 natural amino acids, and then projected to an embedding space. The sequence of embeddings was then fed to the transformer model^28^, consisting of a series of blocks, each composed of a self-attention operation followed by a position-wise multi-layer network (Fig. S1).

Self-attention modules explicitly construct pairwise interactions between all positions in the sequence which enable them to build complex representations that incorporate context from across the sequence. Because the self-attention operation is permutation-equivariant, a positional encoding must be added to the embedding of each token to distinguish its position in the sequence.

### Self-supervised training

Given a large database of protein sequences, a model can be trained using the masked language modelling objective presented in Devlin *et al.^29^*. Each input sequence was corrupted by replacing a fraction of the amino acids with a special mask token. The network was then trained to predict the missing tokens from the corrupted sequence. In practice, for each sequence x, we randomly sampled a set of indices *i* ∈ *M*, for which the amino acid tokens are replaced by a mask token, resulting in a corrupted sequence 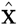. During pre-training, the set M was defined such that 15% of the amino acids in the sequence are corrupted. When corrupted, an amino acid has a 10% chance to be replaced by another randomly selected amino acid and an 80% chance of being masked. During fine-tuning these probabilities do not change, however, only 2.5% of the amino acids in the sequence are corrupted. This probability was selected in order to enable the model to become more accurate for spike protein sequences while keeping its performance on diverse sequences from UniRef100. The training objective corresponds to the negative log-likelihood of the true sequence at the corrupted positions.

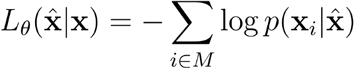

To minimise this loss, the model must learn to identify dependencies between the corrupted and uncorrupted elements of the sequence. Consequently, the learned representations of the proteins, taken as the average of the embeddings of each amino acid (Fig. S.1a), must successfully extract generic features of the biological language of proteins. These features can then be used to fine-tune the model on downstream tasks.

In this work, we used the transformer model from Rives *et al*.^25^ (esm1_t34_670M_UR100) which was trained using the aforementioned procedure on the UniRef100 dataset^47^, containing >277M representative sequences. The pre-trained model was then fine-tuned every month on all the spike protein sequences registered in the GISAID data bank on the specified training date.

Gradient descent was used to minimise the loss function. We relied on the Adam optimiser^48^ and used a learning rate schedule. The batch size was 1. The fine-tuning started with a warm-up period of 100 mini-batches where the learning rate increased linearly from 10^-7^ to 10^-5^. After the warm-up period, the learning rate decreased following 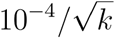 where *k* represents the number of mini-batches.

As the GISAID dataset is imbalanced towards some lineages that have been more prevalent and because certain regions have performed more sequencing than others. To mitigate this bias in the dataset during training, we weighed the importance of each sequence differently in the loss calculation. The importance of a sequence is defined as

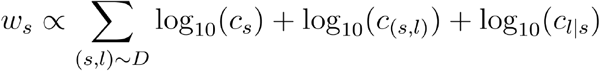

where the values *c_s_* and *c_s,l_* are the numbers of occurrences in the dataset of the sequence s and the sequence-laboratory pair (s, l), respectively. The value *c_l|s_* corresponds to the number of laboratories having reported sequence *s*, which measures the prevalence across regions of the variant.

The model excludes from training all the sequences which have been observed only once in the data set. In this way, it eliminates spurious changes, due to sequencing errors, as well as samples of viruses of subpar evolutionary fitness, which do not spread between patients.

### Inference and machine learning score calculations

Once fine-tuned, the model was used to compute the semantic change and the log-likelihood to characterise a S protein sequence.

Formally, an input sequence was represented by a sequence of tokens defined as x = (*x*_1_,…,*x_n_*) where *n* is the number of tokens and ∀*i* ∈ [1, *n*], 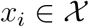 where 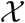 is a finite alphabet that contains the amino acids and other tokens such as class and mask tokens. In this work, a class token was appended to all sequences before feeding them to the network, as such *x*_1_ represents the class token, while *x*_2_,…, *x_n_* represents the amino-acids, or masked amino-acids, in the S protein sequence. The sequence x is passed through attention layers. We note z = (*z*_1_,…, *z_n_*) the output of the last attention layer where *z_i_* is the sequence embedding vector at position *i*.

Please note that in our architecture, the embedding vector *z_i_* is a function of all input tokens (*x_j_*)_*j*∈[1,*n*]_. In opposition, in Bi-LSTM architectures^27^, would be a function of all inputs tokens except the one at the position *i*, (*x_j_*)_*j*∈[1,*n*],*j*≠*i*_.

In order to represent a protein sequence through a single embedding vector, which size does not depend on the protein sequence length, we consider

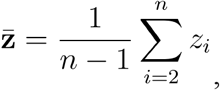

that we call the embedding vector of the variant represented by sequence x. Note that the summation starts at the second position to ensure that the class token’s embedding, which is at the first position, does not contribute to the sequence embedding.

The embeddings of the reference sequences include the Wuhan wild type variants, D614G variants, and other designated variants of concern (Alpha B.1.1.7, Beta B.1.352, Gamma P.1, Delta B.1.617.2, Omicron BA.1, and Omicron BA.2, with the mutations defined in Table S6). In this work, at each date, the semantic change of a variant x was computed with respect to the references with Wuhan variant, D614G variant and variants of concern already designated at the time of detection:

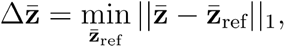

where ||·||_1_ is the L1 norm. This strategy of selecting reference points acts as a proxy for population immunity status due to exposure to proliferating variants.

The last attention layer output z is transformed by a feed-forward layer and a softmax activation into a vector of probabilities over tokens at each position P = (*P*_1_,…,*P_n_*) where *p_i_* is a vector of probabilities at the position *i*, *P_i_* = (*p*(*x_i_* = *x*_1_ | x),…, *p*(*x_i_* = *x_n_* | x)).

The log-likelihood of a variant *l*(x) is computed from these probabilities. It is calculated as the sum of the log probabilities over all the positions of the spike protein amino acids. Formally,

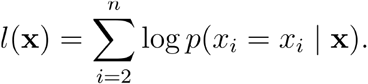

This quantity measures the likelihood of observing a variant sequence x according to the model. Therefore, the more sequences in the training data that are similar to a considered variant, the higher the log-likelihood will be for the variant. The proposed log-likelihood metric supports substitution, insertion, and deletion without the requirement of a reference.

### Implementation details

The method is implemented using the Pytorch^49^ deep learning framework. Model training and inferences were performed on InstaDeep’s high performance computing infrastructure. The average training and inference times were <4 GPU days and <12 GPU hours, respectively, using Nvidia A100-SXM4-40GB GPUs.

### Epitope alteration score

Epitope alteration score attempts to capture the impact of mutations in the variant in question on recognition by experimentally assessed antibodies. Specifically, 332 previously experimentally resolved structures of neutralising antibodies (nAbs)^14–17^ were collected from the coronavirus antibody database on April 22nd, 2022. 719 antibody epitopes were computed for the epitope alteration score calculation.

For each structure, an epitope per antibody was calculated as the set of positions that are in contact with the heavy or light chain of an antibody, where two residues are considered to be in contact if the smallest Euclidean distance between their atoms is smaller than 4 Ångstroms. The epitope can be defined across multiple spike chains. An antibody was considered to be “escaped” by a variant if any position of its epitope is mutated.

Given a variant’s S protein sequence, we first computed the mutation with respect to wild type. We then count the number of *unique* antibodies involving altered positions, as measured across all the antibody-spike complex structures, to estimate how likely the mutated variant is to escape recognition by an antibody.

This score, by design, highly emphasises the effect of mutations on highly antigenic sites, such as the receptor-binding domain (RBD). However, this score is not only limited to the RBD as the epitope calculation pertains to the entire spike protein.

### ACE2 binding score

We selected 402 receptor-binding domain (RBD) differentiated variants, including the wild type, for *in silico* simulation. For each variant, we used an implementation of the ML-based protein complex structure prediction pipeline, analogous to one of Jumper *et al.^50^*, to produce a putative structure of the spike protein RBD-ACE2 complex structure. During the structure generation, standard optimisation methodologies were applied to the coordinates of the predicted complex to optimise the knowledge-based potential, analogous to the methods proposed in Alford *et al. ^51^* This initial estimate was subsequently used for a local complex optimisation protocol, repeated 500 times per each estimate.

In this protocol, we performed the following steps: translate both chains away from each other by 100 Å; minimise the structures with regard to the knowledge-based potential and stored its value as a “separated” energy metric; perform Monte Carlo sampling with fully flexible structures aiming to minimise the total energy of the complex with regard to the knowledge-based potential, which are stored as “complexed” energy metrics; compute the difference between “complexed” and “separated” metrics and denote it as “dG”; measure the difference in solvent accessible surface area between the “separated” and “complexed” structures and denote it as “dSASA”; and finally compute the ratio of “dG” and “dSASA”, which denotes the interaction strength per unit of solvent accessible surface area.

This protocol resulted in >250,000 structures in total, which were used for this work. For each structure, we calculated the change of binding energy when the interface forming chains were separated, versus when they were complexed. These measurements were aggregated per RBD variant using medians. Each metric was then normalised to the corresponding wild type metric. The metrics for wild type are all assigned a value of 1 because the wild type corresponds to no mutation in the RBD.

For sequences having other RBD mutation combinations for which we do not have simulation results, a KNeighborsRegressor model was built where the training data included sequences with known RBDs (for which we have simulation results) along with their language model embedding and energy metrics. Given a sequence with an RBD for which we do not have simulation results, 7 neighbour sequences are identified in the embedding space and their average energy metrics are used as an estimation. The estimated energy metrics are further averaged across all sequences per RBD to ensure that the same RBD mutation combination has the same estimated metrics.

### Growth score

The growth score was computed from the GISAID metadata. At a given date, we considered only sequences that had been submitted within the last eight weeks. For each lineage, its proportion among all submissions was calculated for the eight-week window and for the last week, denoted by r_win_ and r_last_, respectively. The growth of the lineage is defined by their ratio, r_last_ / r_win_, measuring the change of the proportion. Having a value larger than one indicates that the lineage is rising in prevalence and smaller than one indicates a decline.

It is important to note that, due to its nature, growth is only meaningful beyond trivial initial periods with a count in a range of at least tens of observations. To eliminate the trivial spikes in growth (e.g., a threefold change from a count of 1 to count of 3), the EWS computed this metric with regard to the total count of the base lineage (as assigned by PANGOLIN ^44^) and not the individual sequence.

### Score scaling and merging

The semantic change, log-likelihood, epitope alteration score, ACE2 binding score, and growth have different scales and units. Thus, they can not be compared directly. To make comparisons possible, a scaling strategy was introduced. For a given metric, m, all the variants considered are ranked according to this metric. In the ranking system used, the higher the rank the better. Variants with the same value will get the same rank. The ranks are then transformed into values between 0 and 100 through a linear projection to obtain the values for the scaled metric. All reported scores, except for log-likelihood, have been scaled according to this strategy.

Log-likelihood was observed to strongly penalise variants with a large number of mutations. An increased number of mutations may strongly affect fitness, thus explaining decreased log-likelihood. However, as the variants that are scored by the EWS have been registered, which implies that they managed to infect hosts and replicate sufficiently to be detected, we hypothesise that they have at least minimal fitness. A variant with two mutations, whose log-likelihood is in the bottom 20^th^ percentile globally, is less likely to survive the evolutionary competition. A variant, with analogous log-likelihood, but with twenty mutations is more likely among other, similarly mutated variants. Thus, we introduced a group-based ranking strategy where each variant was ranked only among variants with a similar number of mutations. For each variant, having N mutations, its conditional log-likelihood score is ranked among all variants having at least M mutations, with M = min(N, 50). Deletions at N-terminal or C-terminal positions were considered as one single mutation for this grouping. In each group, the ranks were then transformed into values between 0 and 100 through a linear projection to obtain the values for the scaled metric. Although this work uses all the samples that have no less than ten mutations fewer than the query, the results are largely robust to the choice of a threshold.

The immune escape score was computed as the average of the scaled semantic change and of the scaled epitope alteration score. The fitness prior score was computed as the average of the scaled conditional log-likelihood, the scaled ACE2 binding score, and the scaled growth.

### Semantic change vs epitope alteration score

Semantic change is a measure of *how different* is the variant in question with regard to the underlying statistical model (large ML model fine-tuned on spike protein sequences observed until a given time point). This value depends on the observations. Epitope alteration score is a measure of *how many distinct epitopes are evaded* by the variant in question in comparison to wild type. This score, on the other hand, is computed purely based on known binding sites of antibodies, as reported in Protein Data Bank. It too changes with time with new discoveries of anti-spike antibodies, but to a lesser extent and is expected to converge to a stable value.

We observe that these scores, while intuitively correlated, are not collinear. In particular, we see that most of HRVs regarded as immune escaping (and denoted as VoCs, VoIs etc.) indeed exhibit high semantic change, but are rather diverse in terms of Epitope Alteration Score (Fig. S.11).

### Retrospective detection analysis

The retrospective analysis was conducted for each week between September 16th, 2020 and May 15th, 2022 using the same detection protocol. For each week, the language model was trained on variants up to that week. The structures used for epitope alteration score were also limited to those available prior to the analysis date. For the reference sequences of the semantic change score, a variant of concern was included as reference only for the subsequent weeks. Moreover, as the lineage assignments vary through time depending on the PANGOLIN versions, during the analysis, for each sequence, only its associated lineage at the time of analysis was considered. Each sequence was considered available only at the date of its submission and not at the date of collection.

Each week, all sequences reported in the previous eight weeks were ranked. Afterwards, as the objective was to detect variants that are potentially dangerous in the future, each week, we only considered the novel sequences that have never been submitted prior to the current week. Moreover, we exclude the sequences that were labelled as variants of concern at that time. In order to prevent selecting sequences that are highly similar to each other, the candidate sequences were clustered by their RBD mutations. Then for each RBD cluster, only the sequence having the highest score was kept. At last, among all the sequences remaining, the top scoring sequences were selected to make up the watch list.

### Enrichment score

For each week, the enrichment score was calculated as the number of detected HRV sequences by a method divided by the expected number of detected HRV sequences by random sampling. For instance, if there are in total 1,000 sequences as detection candidates, among which 50 sequences (5%) would later be confirmed as HRVs, then with a capacity of 12 sequences, we expect to detect 0.6 sequences (5% of 12) by random sampling. If the method detects 5 sequences, the enrichment is therefore 5 / 0.6 = 8.33.

### Detecting HRVs by standard ML methods

For all the baselines, we considered a protein sequence represented by a vector of **N** binary components. To compute the representation for a protein sequence **S** deposited at time **t**, we calculated the **N** most prevalent mutations in all deposited sequences in the preceding 8 weeks. Each binary component of the representation equals **1** if the mutation is present in **S** and **0** otherwise. We consider **N**=1,280 for all the baselines, to permit a direct comparison with the methods proposed in the paper.

As our method learns from unlabeled data, we first considered the unsupervised learning baseline: Uniform Manifold Approximation and Projection (UMAP). This is an intuitive approach, as it has been successfully applied to analogous problems in biology, and it is known to render meaningful insights when applied in life science settings^52^. UMAP was performed each week over the representations of all sequences available up to the relevant week. A metric equivalent to the semantic change was computed as a min L1 distance between the sequence projection and the projections of reference sequences in UMAP spaces. The same detection technique as performed by the EWS was then used to flag a set of 12 variants suspected to be dangerous each week. Only 9 out of 16 variants were detected using this technique (Table S11). In comparison, analogous techniques applied in the EWS detect all 16 variants, with a mean lead time of 59 days. Both results highlight the need for more involved representations, such as the ones learned by transformer models, especially in tasks like this one where the significance of the novel findings is difficult to approximate *a priori*.

Second, a supervised learning baseline was considered. Each week, we labelled all protein sequences that had been registered with **1** if it had been designated as a HRV anytime before or during the week considered and **0** otherwise. A Generalised Linear Model (GLM) was built over the same previously described 1280-dimensional representation of the sequences. The probability of belonging to the HRV class returned by the GLM was then used to rank sequences. Subsequently, the same detection technique performed by the EWS was used to flag a set of 12 variants suspected to be dangerous each week. We found that 8 of 16 variants were eventually detected before WHO designation (Table S11). In addition, this approach cannot be used early in a pandemic as there are no labels available and hallmark mutations are unlikely to be among the most common ones in the population early on.

## Acknowledgements

Supported by BioNTech and InstaDeep. We thank the BioNTech German clinical trial (NCT04380701, EudraCT: 2020-001038-36) participants, from whom the post-immunisation human sera for the cross-neutralisation analysis were obtained.

## Author contributions

U.S., K.B., M.J.S., A.M., Y.F, T.P. and A.P. conceived and conceptualised the work. K.B. conceived the machine learning scoring models. K.B., Y.F., T.P., and A.L. conceived the machine learning training procedure. K.B., M.J.S., Y.F., N.L.C., A.K, A.L., and I.K. conceived and developed the data pipeline, software, and visuals. Y.F., T.P., and I.K. performed the machine learning experiments. M.J.S. conceived and developed the *in silico* epitope alteration score and structural bioinformatics methodology. A.M. and B.G.L. planned and supervised the *in vitro* experiments. A.M., B.G.L. and B.S. performed *in vitro* experiments. A.M., B.G.L. and B.S. analyzed *in vitro* experimental data. U.S., K.B., M.J.S., Y.F., T.P., N.L.C., A.M., A.U.L., A.P., and Y.L. interpreted data and wrote the manuscript. All authors supported the review of the manuscript.

## Material availability

Biological materials are available from the authors under a material transfer agreement with BioNTech.

