## Supplementary Materials for "Early Computational Detection of Potential High Risk SARS-CoV-2 Variants"

### Supplementary Information

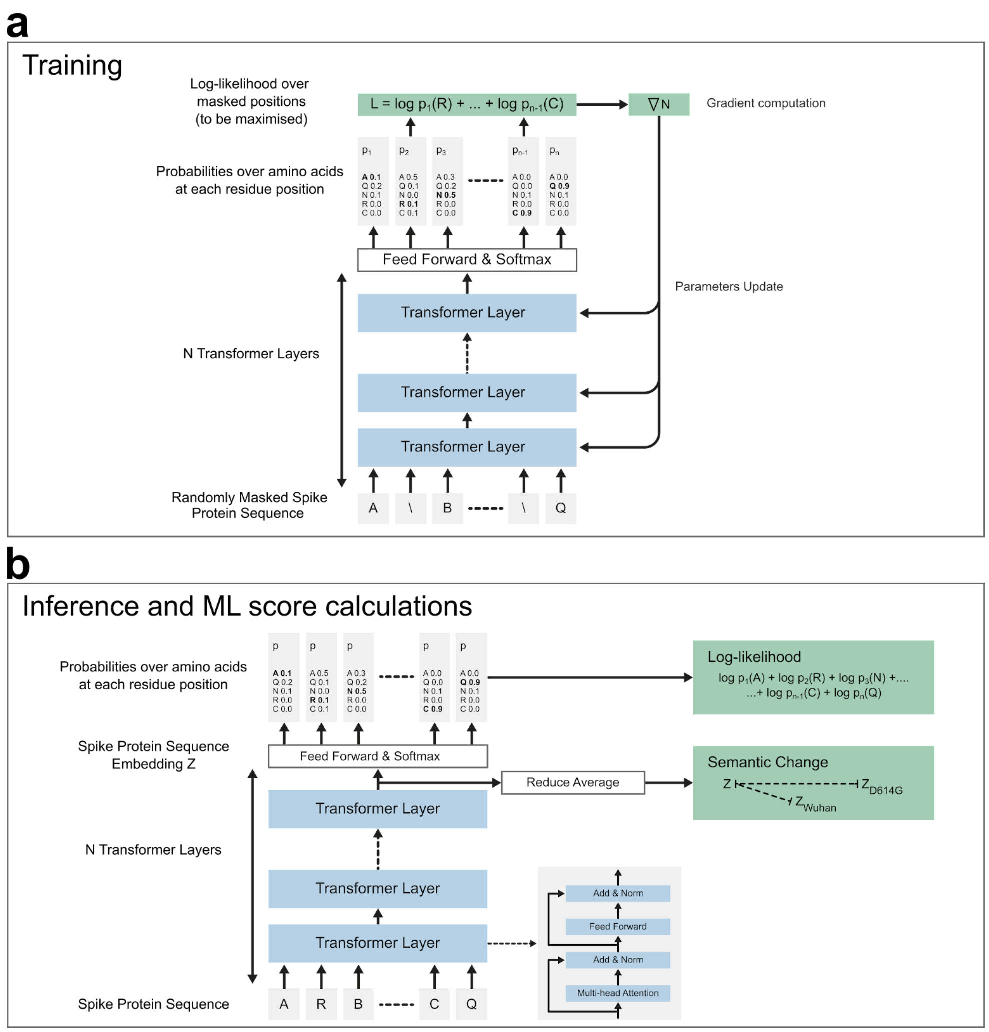


**Figure S1. Machine learning modelling. (a)** A transformer language model is pre-trained on all the protein sequences registered in the UniRef100 dataset. Every week, the model is fine-tuned over all the S protein sequences registered at least twice, so far by the GISAID initiative. Both the pre-training and fine-tuning use the same protocol. Amino acids of a protein sequence are randomly masked. The model predicts probabilities over amino acids at each residue position, both for residues that were masked and not masked. A loss function evaluates the sum over the masked residues of the negative log-probability of the correct predictions. A gradient of this loss is computed and used to update the model's parameters so as to minimise the loss function. **(b)** Once fine-tuned, the model is used to compute the semantic change and the log-likelihood to characterise an S protein sequence. The output of the last transformer layer is averaged over the residues to obtain an embedding z of the protein sequence. The embedding of the references including the Wuhan variant $z_{wuhan}$ and D614G variant $z_{D614G}$ are computed once for all. The semantic change is computed as the minimum L1 distance between the z and reference sequences. The log-likelihood is computed from the probabilities over the residues returned by the model. It is calculated as the sum of the log-probabilities over all the positions of the S protein amino acids.

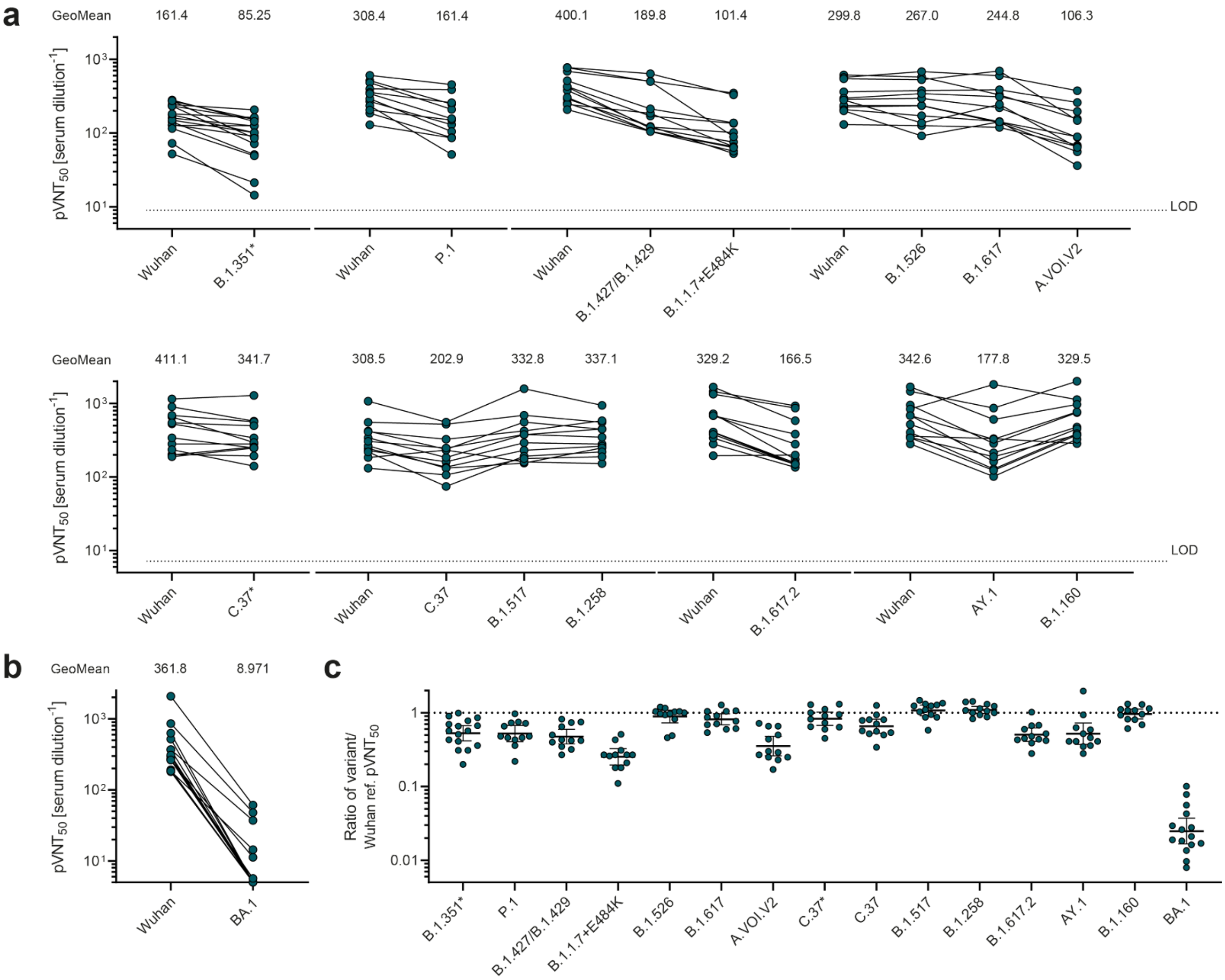


**Figure S2. Cross-neutralisation of BNT162b2-immune sera against VSV-SARS-CoV-2-S pseudoviruses bearing the S protein of selected SARS-CoV-2 variants.** Serum samples were obtained from participants in the BNT162b2 vaccine phase-I/II trial on day 28 or day 43 (7 or 21 days after Dose 2). A recombinant vesicular stomatitis virus (VSV)-based SARS-CoV-2 pseudovirus neutralisation assay was used to measure neutralisation. The pseudoviruses tested incorporated the ancestral SARS-CoV-2 Wuhan Hu-1 spike or spikes with substitutions present in B.1.1.7+E484K (Alpha), B.1.351 (Beta), P.1 (Gamma), B.1.617.2 (Delta), AY.1 (Delta), B.1.427/B.1.429 (Epsilon), B.1.526 (Iota), B.1.617 (Kappa), C.37 (Lambda), C.37* (Lambda), A.VOI.V2, B.1.517, B.1.258, B.1.160, and BA.1 (Omicron) (Table S.5). **(a)** Pseudovirus 50% neutralisation titers ($pVNT_{50}$) are shown. Dots represent results from individual serum samples. Lines connect paired neutralisation analyses performed within one experiment. In total 8 experiments were performed covering the listed SARS-CoV-2 variants always referencing variant-specific neutralisation to the Wuhan reference. **(b)** $pVNT_{50}$ against BA.1 (Omicron) is shown. Dots represent results from individual serum samples. Lines connect paired neutralisation analyses performed within one experiment. **(c)** The ratio of $pVNT_{50}$ between the SARS-CoV-2 variant and Wuhan reference strain spike-pseudotyped VSV. Dots represent results from individual serum samples. Horizontal bars represent geometric mean ratios, error bars represent 95% confidence intervals.

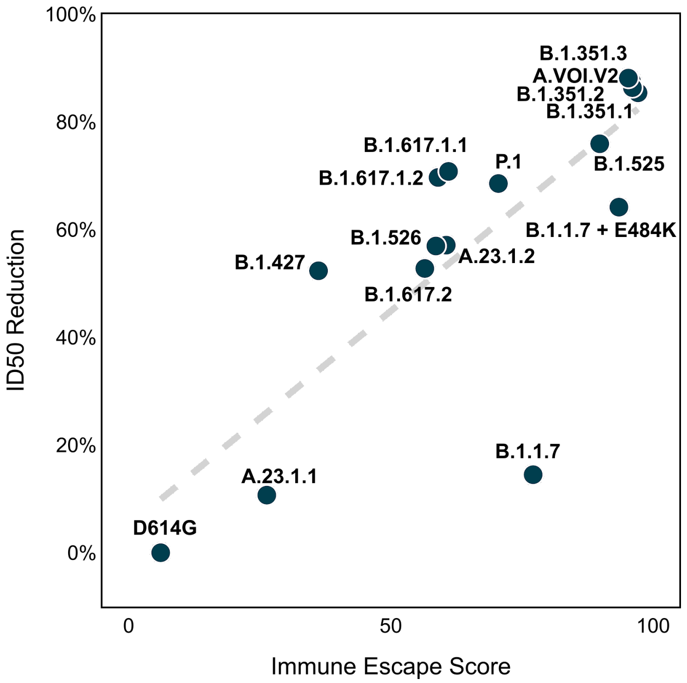


**Figure S3. Validation of immune escape score with serum neutralising activity of mRNA-1273**[1] The regression plot between reduction in immune response (as measured by inhibitory dilution, $ID_{50}$) versus immune escape score across 16 variants. Dashed lines represent the fitted ordinary least squares regression line. Pearson correlation is 0.79 with a *p*-value of 3E-4. Spearman correlation is 0.8 with a *p*-value of 6E-5.

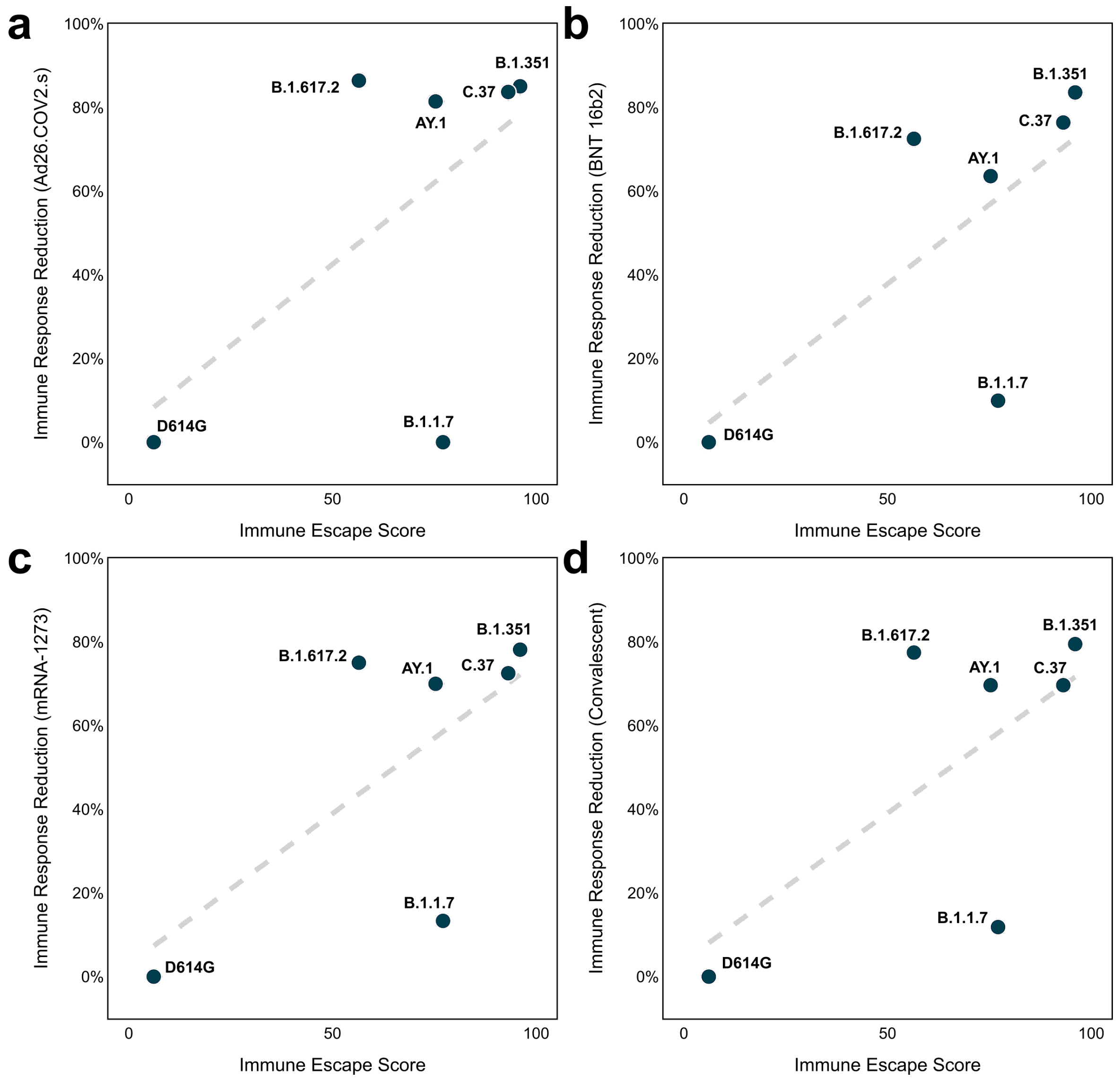


**Figure S4. Validation of immune escape score with neutralising antibody titers[2].** The regression plot between Immune Escape Score and immune response reduction is displayed across 6 variants for (a) Neutralisation by Ad26.COV2.S vaccine-elicited antibodies. Dashed lines represent the fitted ordinary least squares regression line. Pearson correlation is 0.60 with a $p$-value equal to 2E-1. Spearman correlation is 0.29 with a $p$-value equal to 6E-1. (b) Neutralisation by BNT162b2 vaccine-elicited antibodies. Pearson correlation is 0.69 with a $p$-value equal to 1E-1. Spearman correlation is 0.77 with a $p$-value equal to 7E-2. (c) Neutralisation by mRNA-1273 vaccine-elicited antibodies. Pearson correlation is 0.68 with a $p$-value equal to 1E-1. Spearman correlation is 0.60 with a $p$-value equal to 2E-1. (d) Neutralisation by convalescent sera. Pearson correlation is 0.66 with a $p$-value equal to 2E-1. Spearman correlation is 0.55 with a $p$-value equal to 3E-1.

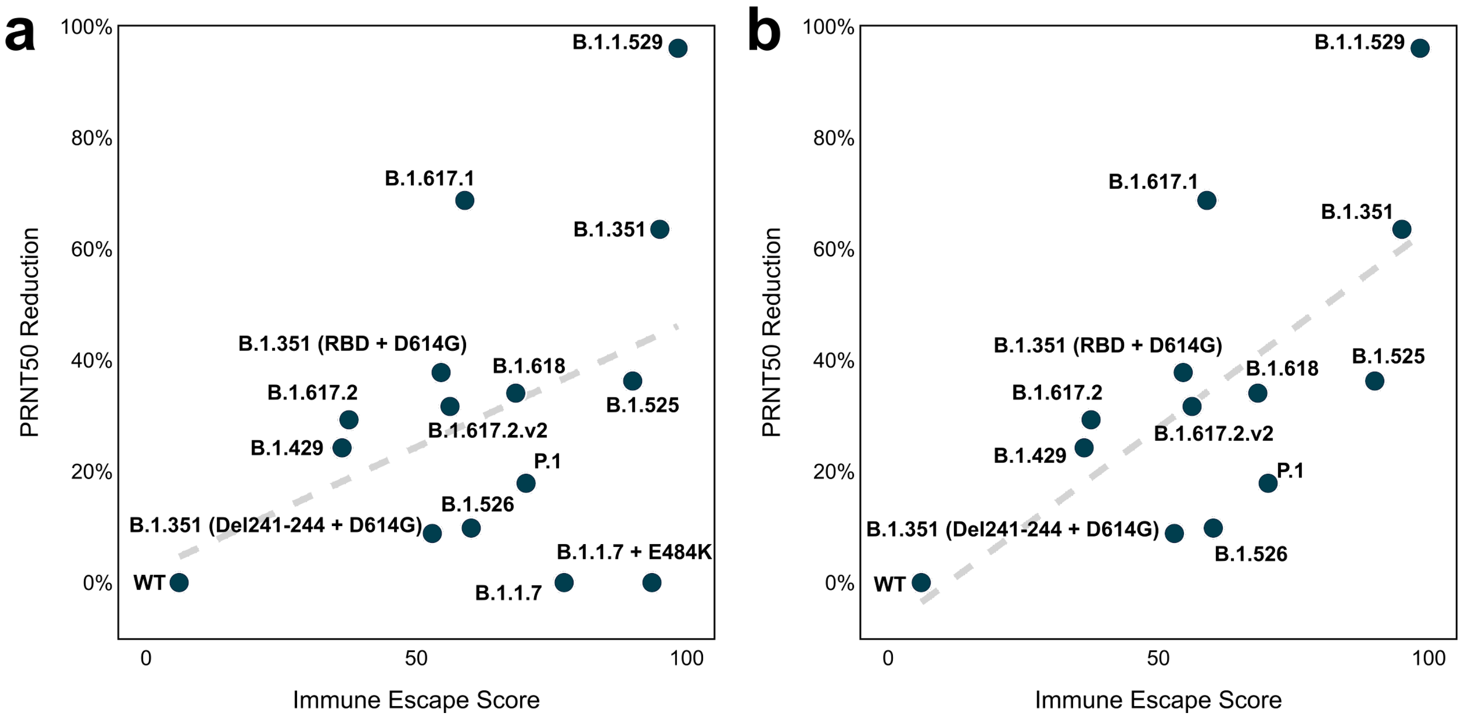


**Figure S5. Validation of immune escape score with neutralising activity of BNT162b2-elicited serum**[3–6] (a) The regression plot between Immune Escape Score and immune response reduction is displayed across 14 variants. The immune response reductions of B.1.1.7 and B.1.1.7+E484K were negative and have been modified to zero. Dashed lines represent the fitted ordinary least squares regression line. Pearson correlation is 0.41 with a *p*-value equal to 1E-1. Spearman correlation is 0.30 with a *p*-value equal to 3E-1. (b) The regression plot between Immune Escape Score and immune response reduction is displayed across 12 variants, without B.1.1.7 and B.1.1.7+E484K. Pearson correlation is 0.68 with a *p*-value equal to 1E-2. Spearman correlation is 0.62 with a *p*-value equal to 3E-2.

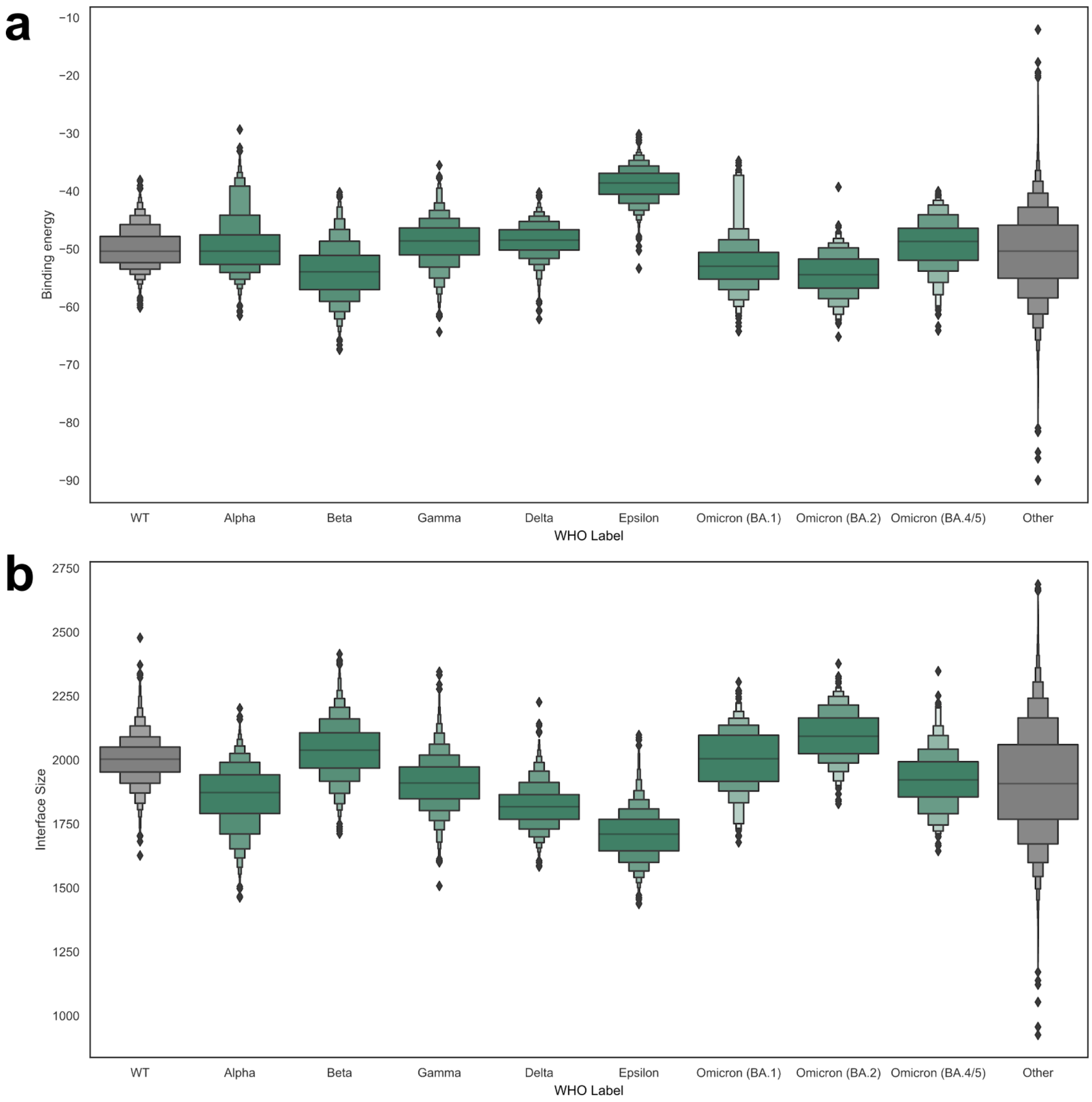


**Figure S6. Results of molecular simulations of RBD binding.** The efficiency of S protein RBD binding to the ACE2 receptor is dictated by the combination of binding energy (a), where a lower value is indicative of better binding, and size of the interface (b). Both box plots depict the distribution of these values across performed RBD binding simulations for circulating S protein variants. Note that, while larger interfaces may be more difficult to form, they are also more difficult to break. Strikingly, Omicron, despite its heavily mutated RBD, has a relatively large interface and a binding affinity around the 75th percentile of the background distribution ('Other') for BA.1 and BA.2, and between 25th and 50th percentile for BA.4 and BA.5.

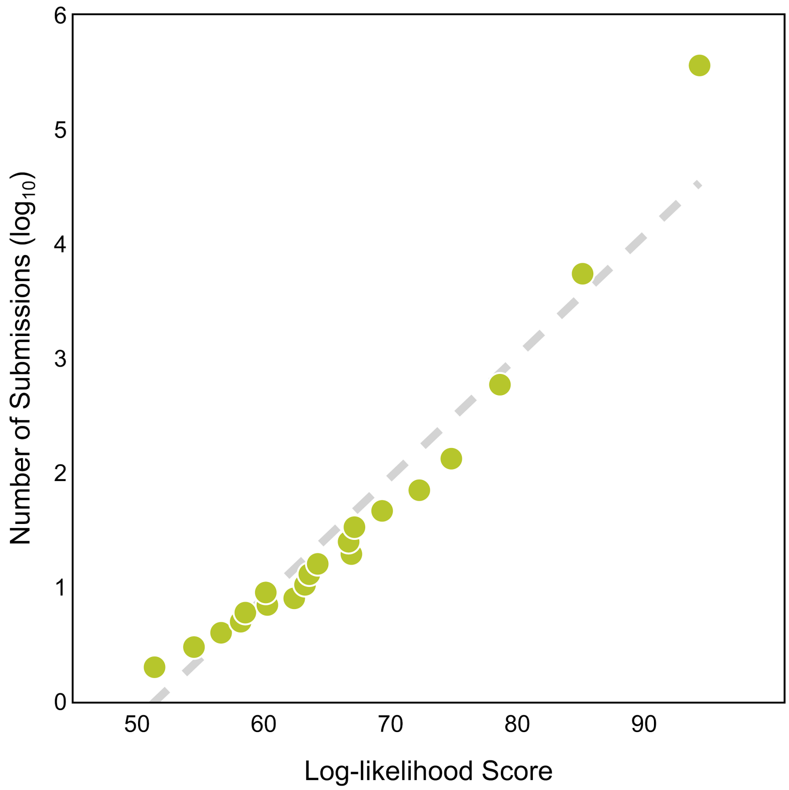


**Figure S7. Validation of the log-likelihood score.** Sequences are grouped into bins based on their submission count and the log-likelihood scores and number of submissions were averaged per bin. The first ten bins correspond to count 1 to 10. The next 10 bins are equally split between counts 11 and 1000 such that each bin has a similar number of sequences. The last two bin contains all sequences having a submission count from 1,000 to 10,000 and sequences having more than 10,000 submissions. The regression plot displays the correlation between the number of submissions and conditional log-likelihood score. Dashed line represents the fitted ordinary least squares regression line. Pearson correlation is 0.96 with a $p$-value equal to 1E-11. Spearman correlation is 0.99 with a $p$-value equal to 3E-20. This shows that the mean conditional log-likelihood of sequences that are observed frequently in circulation is much higher than that of outlier, infrequent sequences.

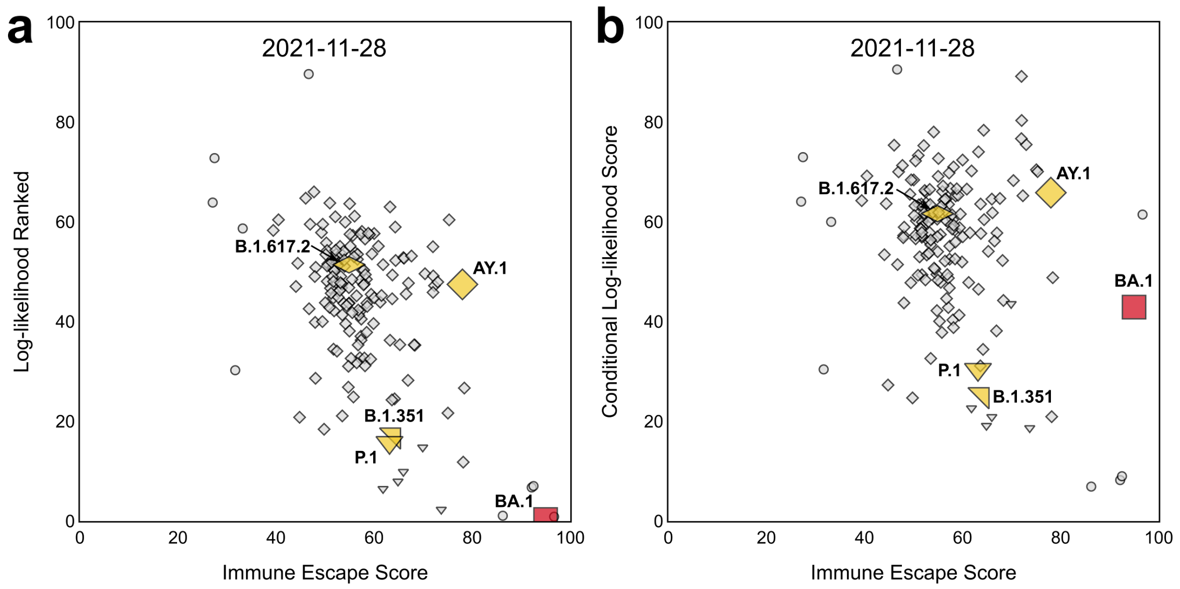


**Figure S8. Log-likelihood score corrects for large mutation count. (a)** Snapshot of lineages in terms of log-likelihood ranked on November 28th 2021. **(b)** The same plot as in (a) but the log-likelihood score has been corrected to account for large mutation count and immune escape score. Red markers indicate the designated lineages of the week, yellow markers are the previously designated lineages and grey markers indicate other lineages. Small circles correspond to non-VOC lineages and other symbols correspond to designated variants and their closely related lineages (e.g., squares correspond to Omicron BA.1 and its closely related lineages). Only lineages that have been observed within the past 8 weeks and have been reported more than 10 times are included.

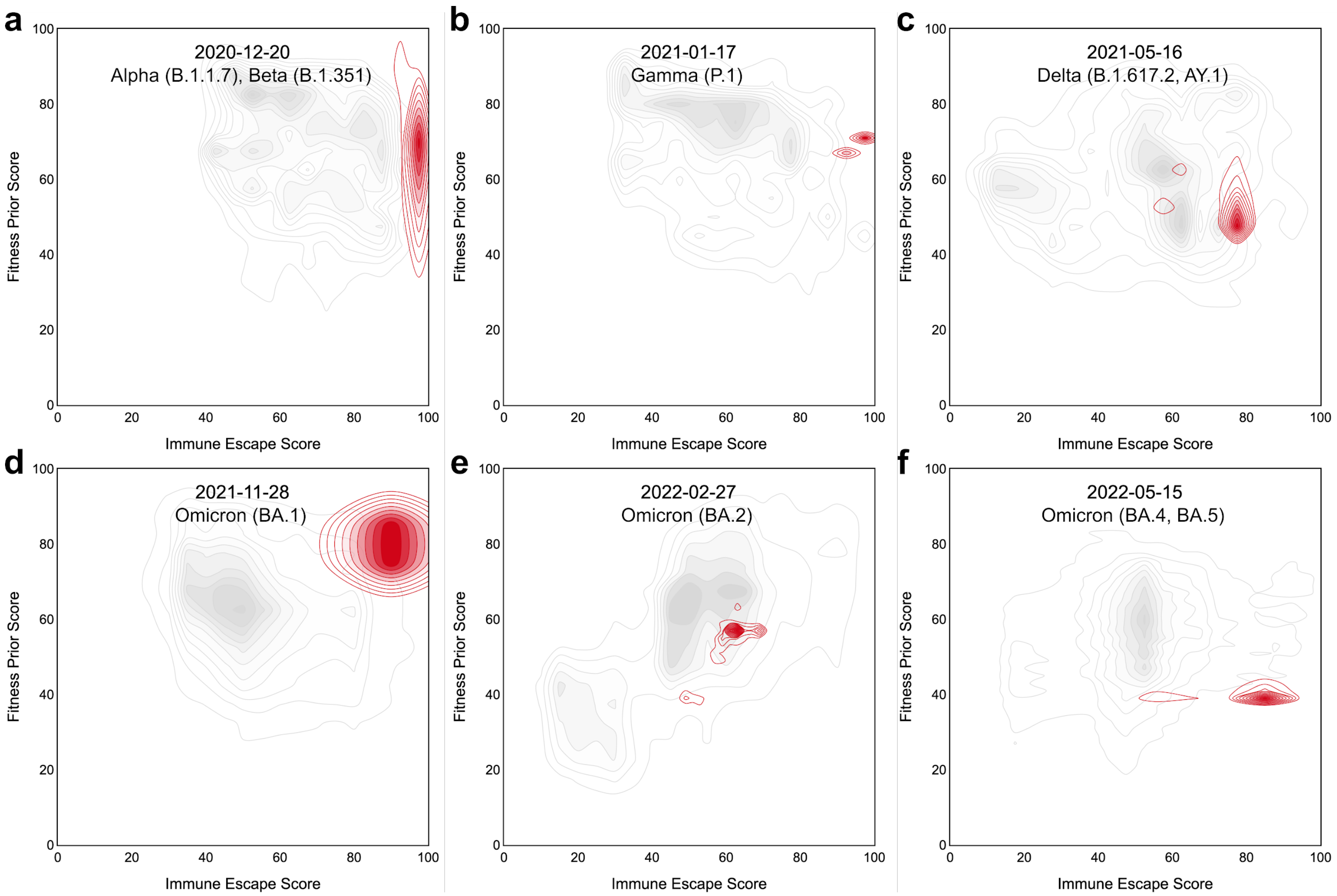


**Figure S9. Combining immune escape and fitness prior for continuous monitoring.** Density contour plots of sequences in terms of fitness prior and immune escape score on (a) December 20th 2020, (b) January 17th 2021, (c) May 16th 2021, (d) November 28th 2021, (e) February 27th 2022, and (f) May 15th 2022, corresponding to the week of the designations of Alpha/Beta, Gamma, Delta, Omicron BA.1, Omicron BA.2 and Omicron BA.4/BA.5. Red contours indicate the designated lineages (also indicated on each plot) of the week and other lineages' markers are grey. Only sequences of the lineages that have been observed within the past 8 weeks and have been reported more than 10 times are included. Note how Alpha, Beta, Gamma, and BA.1 emerged with a very clear signal (see panel a, b, and d respectively). Omicron BA.4 and BA.5 appeared as highly immune escaping, yet relatively fit variants (see panel f), while initially reported samples of Delta and BA.2 were not immediately striking as alarming (see panels c and e), however, their subsequent reports exhibited much more alarming metrics, permitting for their timely detection.

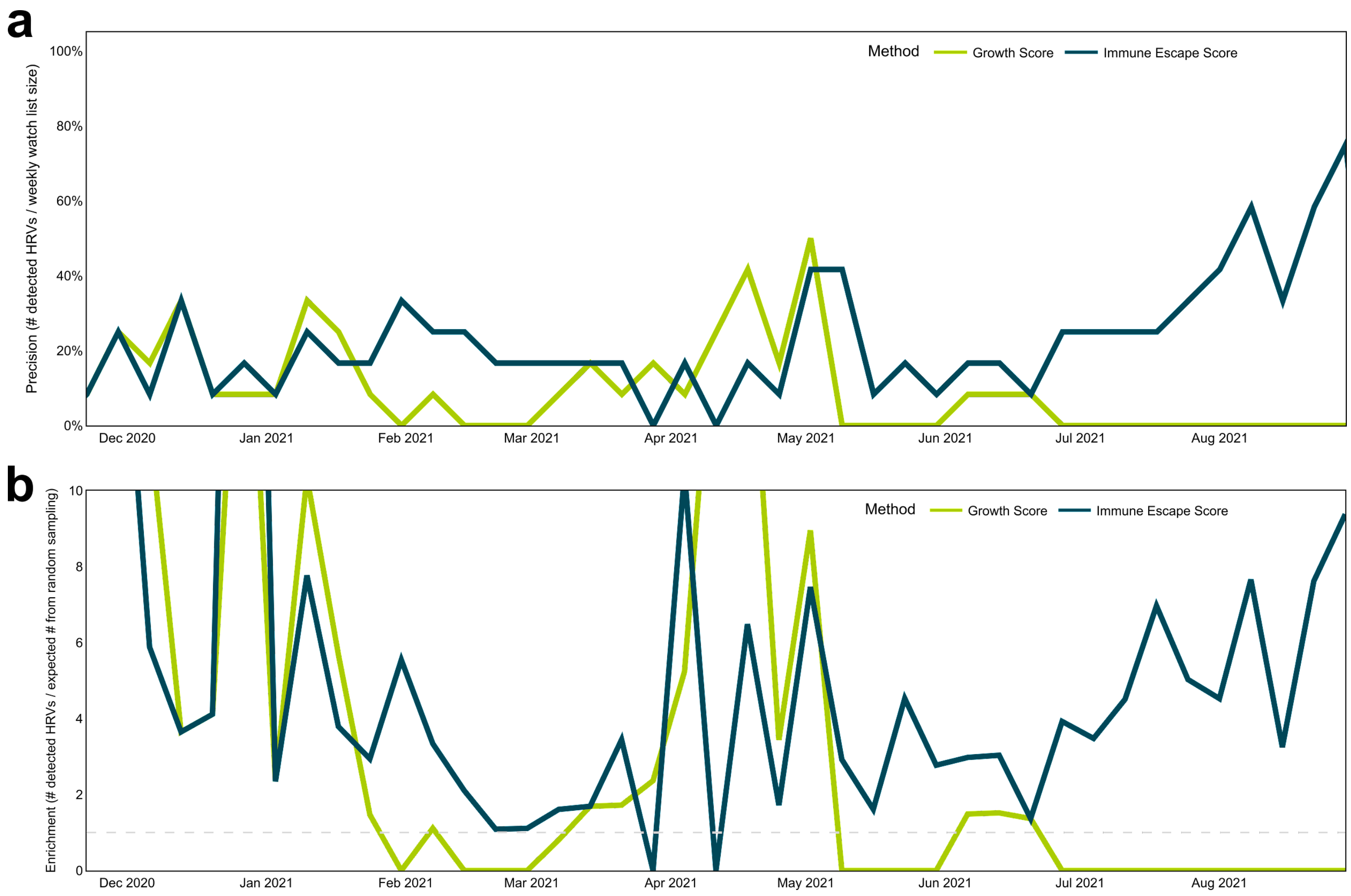


**Figure S10. Precision and Enrichment Analysis.** For each week, the precision (the number of EWS detected HRVs divided by the weekly watch list size) and enrichment (the number of EWS detected HRVs divided by the expected number of random sampling detected HRVs) were calculated from July 2021 to October 2021. For enrichment, the dash line is enrichment equals one. Across the weeks, the immune escape score achieved a significantly better enrichment when there were many new variants growing in prevalence. In particular observe the performance of the EWS system in situations when multiple, viable lineages are simultaneously in circulation (February-March 2021 and May 2021 onwards), which permits for detection of *new* threats, such as dangerous sublineages (such as AY.* and Lambda, given globally circulating B.1.617.2).

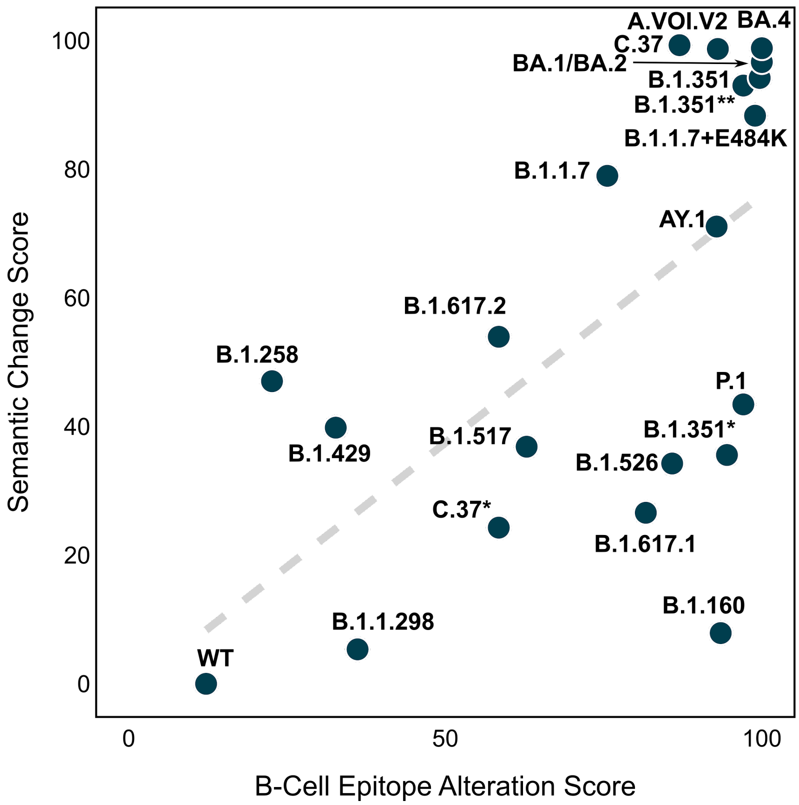


**Figure S11. Metrics of anticipated reduction of the immune response.** Semantic change and epitope alteration score accurately segment the variant landscape, allowing for discrimination between variants that do not have immune escape propensity (B.1.429, WT), highly mutated, but neutralisable variants (P.1, B.1.160), and those with high potential for evading immune response (B.1.1.7, AY.1, B.1.351). The dashed line represents the fitted ordinary least squares regression line. Pearson correlation is 0.62 with a *p*-value equal to 2E-3. Spearman correlation is 0.63 with a *p*-value equal to 2E-3.

#### Table S1. N=332 nAbs Resolved 3D Structures PDB Identifiers.

| Code | Last Updated Date | Code | Last Updated Date | Code | Last Updated Date | Code | Last Updated Date | Code | Last Updated Date |
|---|---|---|---|---|---|---|---|---|---|
| 7BZ5 | 2020-05-20 | 6ZLR | 2021-05-05 | 7D6I | 2021-10-18 | 7WD8 | 2022-01-31 | 7PRZ | 2022-01-31 |
| 7K8W | 2020-06-01 | 7M6D | 2021-05-05 | 7MZK | 2021-10-18 | 7QNY | 2022-01-31 | 7OR9 | 2022-01-31 |
| 7BYR | 2020-06-10 | 7A5R | 2021-05-05 | 7MZF | 2021-10-18 | 7ND5 | 2022-01-31 | 7NXB | 2022-01-31 |
| 7K9K | 2020-06-27 | 7E23 | 2021-05-07 | 7MZG | 2021-10-18 | 7QNW | 2022-01-31 | 7NXA | 2022-01-31 |
| 7K9J | 2020-06-27 | 7M6F | 2021-05-07 | 7MZH | 2021-10-18 | 7L2F | 2022-01-31 | 7NX9 | 2022-01-31 |
| 6XEY | 2020-07-26 | 7M6H | 2021-05-07 | 7S0C | 2021-10-18 | 7EZV | 2022-01-31 | 7NX8 | 2022-01-31 |
| 7JMP | 2020-07-31 | 7M6E | 2021-05-07 | 7K9H | 2021-10-18 | 7B3O | 2022-01-31 | 7NX7 | 2022-01-31 |
| 7JMO | 2020-07-31 | 7M6G | 2021-05-07 | 7K9I | 2021-10-18 | 7LSS | 2022-01-31 | 6XC2 | 2022-01-31 |
| 6XKQ | 2020-08-27 | 7DET | 2021-05-11 | 7MZI | 2021-10-18 | 7N5H | 2022-01-31 | 6XC3 | 2022-01-31 |
| 7CHB | 2020-10-01 | 7MF1 | 2021-05-11 | 7V2A | 2021-11-01 | 7NEH | 2022-01-31 | 7C01 | 2022-03-04 |
| 7CH5 | 2020-10-01 | 7DEU | 2021-05-11 | 7V26 | 2021-11-01 | 7NEG | 2022-01-31 | 7E8M | 2022-03-04 |
| 7DCX | 2020-12-14 | 7DEO | 2021-05-11 | 7CWS | 2021-12-02 | 7BEM | 2022-01-31 | 7MJL | 2022-03-04 |
| 7DD8 | 2020-12-14 | 7MKM | 2021-05-15 | 7D4G | 2021-12-02 | 7ND4 | 2022-01-31 | 7MJJ | 2022-03-04 |
| 7DCC | 2020-12-14 | 7MKL | 2021-05-15 | 7N4L | 2021-12-02 | 7ND6 | 2022-01-31 | 7MMO | 2022-03-04 |
| 7DK7 | 2020-12-14 | 7M7B | 2021-05-27 | 7CWO | 2021-12-02 | 7ND8 | 2022-01-31 | 7MJK | 2022-03-04 |
| 7DK4 | 2020-12-14 | 7CJF | 2021-05-27 | 7CWM | 2021-12-02 | 7ND7 | 2022-01-31 | 7THT | 2022-03-04 |
| 7DK6 | 2020-12-14 | 7M8J | 2021-05-27 | 7CWT | 2021-12-02 | 7BEH | 2022-01-31 | 7SOD | 2022-03-04 |
| 7DK5 | 2020-12-14 | 7M71 | 2021-05-27 | 7N4J | 2021-12-02 | 7NDA | 2022-01-31 | 7VYR | 2022-03-04 |
| 7KFY | 2020-12-14 | 7KQB | 2021-05-27 | 7N4I | 2021-12-02 | 7ND9 | 2022-01-31 | 7TP4 | 2022-03-04 |
| 7KFX | 2020-12-14 | 7KQE | 2021-05-27 | 7CWL | 2021-12-02 | 7NDB | 2022-01-31 | 7TP3 | 2022-03-04 |
| 7DD2 | 2020-12-14 | 7AKD | 2021-05-27 | 7CYH | 2021-12-02 | 7NDD | 2022-01-31 | 7WCR | 2022-03-04 |
| 7KZB | 2020-12-21 | 7DZX | 2021-06-06 | 7KLH | 2021-12-02 | 7NDC | 2022-01-31 | 7WCZ | 2022-03-04 |
| 7L02 | 2021-01-05 | 7DZY | 2021-06-06 | 7KMK | 2021-12-02 | 7BEK | 2022-01-31 | 7WD0 | 2022-03-04 |
| 7L06 | 2021-01-05 | 7M6I | 2021-06-10 | 7KML | 2021-12-02 | 7BEJ | 2022-01-31 | 7WD7 | 2022-03-04 |
| 7L09 | 2021-01-05 | 7DJZ | 2021-06-16 | 7KXK | 2021-12-02 | 7CHH | 2022-01-31 | 7PR0 | 2022-03-04 |
| 7L2C | 2021-01-12 | 7DK0 | 2021-06-16 | 7KXJ | 2021-12-02 | 7BEI | 2022-01-31 | 7PQZ | 2022-03-04 |
| 7LQW | 2021-01-12 | 7CH4 | 2021-06-16 | 7KLG | 2021-12-02 | 7MSQ | 2022-01-31 | 7F7E | 2022-03-04 |
| 7KMI | 2021-01-30 | 7E86 | 2021-06-16 | 7CYP | 2021-12-02 | 7QNX | 2022-01-31 | 7L2E | 2022-03-04 |
| 7KMH | 2021-01-30 | 7E7Y | 2021-06-16 | 7CWU | 2021-12-02 | 7CHE | 2022-01-31 | 7Q0G | 2022-03-04 |
| 7LCN | 2021-01-30 | 7E7X | 2021-06-16 | 7ORB | 2021-12-28 | 6XC7 | 2022-01-31 | 7SC1 | 2022-03-04 |
| 7K8X | 2021-01-30 | 7E88 | 2021-06-16 | 7BEL | 2021-12-28 | 7CHC | 2022-01-31 | 7Q0A | 2022-03-04 |
| 7K8Y | 2021-01-30 | 7BEO | 2021-07-09 | 7M3I | 2021-12-28 | 7CHF | 2022-01-31 | 7LD1 | 2022-03-04 |
| 7K8M | 2021-02-12 | 7NTC | 2021-07-09 | 7R7N | 2021-12-28 | 6ZDG | 2022-01-31 | 7PQY | 2022-03-04 |
| 7K8S | 2021-02-12 | 7CHO | 2021-07-24 | 7JV6 | 2021-12-28 | 6ZCZ | 2022-01-31 | 6WPS | 2022-03-04 |

| | | | | | | | | | |
|---|---|---|---|---|---|---|---|---|---|
| 7K8U | 2021-02-12 | 7CHS | 2021-07-24 | 7LQV | 2021-12-28 | 7RW2 | 2022-01-31 | 7THE | 2022-03-04 |
| 7K8T | 2021-02-12 | 7CHP | 2021-07-24 | 7LXW | 2021-12-28 | 7K8V | 2022-01-31 | 7JW0 | 2022-03-04 |
| 6XKP | 2021-02-18 | 7E5Y | 2021-08-05 | 7SO9 | 2021-12-28 | 7K45 | 2022-01-31 | 7L0N | 2022-03-04 |
| 7L57 | 2021-02-28 | 7F62 | 2021-08-16 | 7N8I | 2021-12-28 | 7K4N | 2022-01-31 | 7SOC | 2022-03-04 |
| 6XDG | 2021-02-28 | 7R8M | 2021-08-16 | 7N8H | 2021-12-28 | 7LRT | 2022-01-31 | 7TLY | 2022-03-04 |
| 7RAL | 2021-02-28 | 7R8N | 2021-08-16 | 7KS9 | 2021-12-28 | 7LRS | 2022-01-31 | 7R6X | 2022-03-04 |
| 7DPM | 2021-02-28 | 7R8O | 2021-08-16 | 7SN2 | 2021-12-28 | 7CM4 | 2022-01-31 | 7SOB | 2022-03-04 |
| 7RA8 | 2021-02-28 | 7EJ4 | 2021-08-16 | 7SN3 | 2021-12-28 | 7MLZ | 2022-01-31 | 7TN0 | 2022-03-04 |
| 7L5B | 2021-02-28 | 7M42 | 2021-09-02 | 7LXY | 2021-12-28 | 7MM0 | 2022-01-31 | 7R6W | 2022-03-04 |
| 7M7W | 2021-02-28 | 7F63 | 2021-09-02 | 7DK2 | 2021-12-28 | 7CAI | 2022-01-31 | 6WPT | 2022-03-04 |
| 7D0B | 2021-03-11 | 7E3B | 2021-09-02 | 7FAF | 2021-12-28 | 7CAK | 2022-01-31 | 7JX3 | 2022-03-04 |
| 7D0C | 2021-03-11 | 7E39 | 2021-09-02 | 7JV4 | 2021-12-28 | 7CWN | 2022-01-31 | 7BEP | 2022-03-04 |
| 7N3I | 2021-03-11 | 7L7E | 2021-09-02 | 7K43 | 2021-12-28 | 7CAC | 2022-01-31 | 7MW2 | 2022-04-20 |
| 7LOP | 2021-03-11 | 7L7D | 2021-09-02 | 7JVA | 2021-12-28 | 7E5S | 2022-01-31 | 7WUH | 2022-04-20 |
| 7LAB | 2021-03-11 | 7EJ5 | 2021-09-02 | 7LY2 | 2021-12-28 | 7K8Z | 2022-01-31 | 7WPD | 2022-04-20 |
| 7R8L | 2021-03-11 | 7EH5 | 2021-09-02 | 7JVC | 2021-12-28 | 7K90 | 2022-01-31 | 7MW6 | 2022-04-20 |
| 7D0D | 2021-03-11 | 7K9Z | 2021-10-18 | 7JV2 | 2021-12-28 | 6ZER | 2022-01-31 | 7MW5 | 2022-04-20 |
| 7LAA | 2021-03-19 | 7RR0 | 2021-10-18 | 7FAE | 2021-12-28 | 6ZFO | 2022-01-31 | 7WRV | 2022-04-20 |
| 7LS9 | 2021-03-19 | 7MZN | 2021-10-18 | 7PRY | 2021-12-28 | 6ZDH | 2022-01-31 | 7WPF | 2022-04-20 |
| 7EAM | 2021-03-19 | 7S0B | 2021-10-18 | 7WKA | 2022-01-31 | 7E5R | 2022-01-31 | 7WUE | 2022-04-20 |
| 7KFW | 2021-03-23 | 7E3C | 2021-10-18 | 7WDF | 2022-01-31 | 7PS6 | 2022-01-31 | 7MW3 | 2022-04-20 |
| 7L2D | 2021-03-25 | 7N64 | 2021-10-18 | 7WD9 | 2022-01-31 | 7PS5 | 2022-01-31 | 7ENG | 2022-04-20 |
| 7LM8 | 2021-04-06 | 7MZM | 2021-10-18 | 7BWJ | 2022-01-31 | 7PS4 | 2022-01-31 | 7WED | 2022-04-20 |
| 7LJR | 2021-04-06 | 7JMW | 2021-10-18 | 7WK8 | 2022-01-31 | 6XC4 | 2022-01-31 | 7ENF | 2022-04-20 |
| 7L56 | 2021-04-19 | 7EY4 | 2021-10-18 | 7WK9 | 2022-01-31 | 7PS2 | 2022-01-31 | 7E9P | 2022-04-20 |
| 7EAN | 2021-04-21 | 7EY5 | 2021-10-18 | 7W9F | 2022-01-31 | 7PS7 | 2022-01-31 | 7F46 | 2022-04-20 |
| 7LXX | 2021-04-21 | 7EYA | 2021-10-18 | 7TAS | 2022-01-31 | 7NX6 | 2022-01-31 | 7E9N | 2022-04-20 |
| 7LXZ | 2021-04-21 | 7E5O | 2021-10-18 | 7TAT | 2022-01-31 | 7Q0H | 2022-01-31 | 7KMG | 2022-04-20 |
| 7LY3 | 2021-04-21 | 7S0E | 2021-10-18 | 7Z0Y | 2022-01-31 | 7PS1 | 2022-01-31 | 7L3N | 2022-04-20 |
| 7LY0 | 2021-04-21 | 7N62 | 2021-10-18 | 7Z0X | 2022-01-31 | 7PS0 | 2022-01-31 | 7T01 | 2022-04-20 |
| 7A5S | 2021-05-05 | 7MZL | 2021-10-18 | 7E8F | 2022-01-31 | 7ORA | 2022-01-31 | 7MW4 | 2022-04-20 |
| 6W41 | 2021-05-05 | 7S0D | 2021-10-18 | 7E8C | 2022-01-31 | 7BEN | 2022-01-31 | 7FJO | 2022-04-20 |
| 6ZH9 | 2021-05-05 | 7MZJ | 2021-10-18 | | | | | | |

**Table S2. EWS Scores.** The EWS has in total five sub-scores grouped into immune escape and fitness prior scores. Each sub-score is normalised ranks that range between 0% and 100%. The average of the sub-scores in each score category is computed to define immune escape and fitness prior scores.

| **Score Name** | **Score Category** | **Description** |
|---|---|---|
| Epitope Alteration | Immune Escape | Measures the alteration of the spike protein at epitope positions by counting the number of antibodies potentially escaped. |
| Semantic Change | Immune Escape | Measures the functional change of the spike protein using Transformer-derived embedding differences with respect to the references including wild type, D614G, and other variants of concern. |
| ACE2 Binding | Fitness Prior | Approximates the binding affinity using binding energies estimated from in-silico simulations. |
| Conditional Log-Likelihood | Fitness Prior | Measures the relative rank of the measured existence probability of the spike protein using Transformer-derived log-likelihoods with reference to other sequences with similar mutation count. |
| Growth | Fitness Prior | Calculated lineage-level growth using deposited sequence metadata (retrospective data). |

**Table S3. Antibody Class Distribution.** Among the 227 antibodies in collected data, 88 have defined antibody class in the data collected from https://jbloomlab.github.io/SARS2_RBD_Ab_escape_maps/.

| Antibody Class | Number of antibodies |
|---|---|
| Class 1 | 38 |
| Class 2 | 20 |
| Class 3 | 19 |
| Class 4 | 11 |

**Table S4. Epitope Class Distribution.** For each of the epitopes defined in Sikora et al.[7], we count the number of relevant epitopes and antibodies. An antibody can have multiple epitopes defined within different structures. Each epitope corresponds to a set of positions and it is assigned to all the classes that contain one of these positions. For instance, the epitope of an antibody interacting with spike protein at positions 164, 332, and 333 is assigned to the epitope class E3 and E4.

| Epitope Class | Positions Range | Number of Epitopes | Number of Antibodies |
|---|---|---|---|
| E1 | 15-28, 63-79, 247-260 | 76 | 20 |
| E2 | 97, 178-189, 207-219 | 14 | 5 |
| E3 | 37-164 | 65 | 19 |
| E4 | 332-346 | 103 | 47 |
| E5 | 403-406, 438, 440-451, 495-506 | 429 | 158 |
| E6 | 452-476, 479-482, 484-494 | 417 | 156 |
| E7 | 527-537 | 18 | 6 |
| E8 | 603-605, 633-624, 656-661, 674-693 | 3 | 1 |
| E9 | 808-814 | 76 | 20 |

**Table S5. Epitope Region Distribution.** For each of the four different regions on spike (S) protein, we count the number of relevant antibody epitopes. An antibody can have multiple epitopes defined within different structures. Each epitope corresponds to a set of positions and it is assigned to all the regions that contain one of these positions. For instance, epitope of an antibody interacting with spike protein at positions 164, 332, and 333 is assigned to NTD, RBD, and S1.

| Region | Positions Range | Number of Epitopes | Number of Antibodies |
|---|---|---|---|
| NTD | 13-303 | 94 | 28 |
| RBD | 319-541 | 632 | 204 |
| S1 | 13-685 | 715 | 226 |
| S2 | 686-1273 | 7 | 2 |

**Table S6. Spike protein mutations in SARS-CoV-2 spike pseudoviruses and observed reduction of neutralising antibody response in pseudovirus neutralisation assay.**

| Lineage | WHO Nomenclature | Mutation | Reduction |
|---|---|---|---|
| B.1.1.7 | Alpha | H69- V70- Y144- N501Y A570D D614G P681H T716I S982A D1118H | 22.92% |
| B.1.1.7+E484K | Alpha | H69- V70- Y144- E484K N501Y A570D D614G P681H T716I S982A D1118H | 74.65% |
| B.1.351 | Beta | L18F D80A D215G L242- A243- L244- R246I K417N E484K N501Y D614G A701V | 80.04% |
| B.1.351* | Beta | D80A D215G L242H K417N E484K N501Y D614G A701V | 47.19% |
| B.1.351** | Beta | D80A D215G L242- A243- L244- K417N E484K N501Y D614G A701V | 77.45% |
| P.1 | Gamma | L18F T20N P26S D138Y R190S K417T E484K N501Y H655Y T1027I V1176F | 47.66% |
| B.1.617.2 | Delta | T19R G142D E156G F157- R158- K417N L452R T478K D614G P681R D950N | 49.43% |
| AY.1 | Delta | T19R T95I G142D E156G F157- R158- W258L K417N L452R T478K K558N D614G P681R D950N | 48.09% |
| B.1.427/B.1.429 | Epsilon | S13I W152C L452R D614G | 52.57% |
| B.1.526 | Iota | L5F T95I D253G E484K D614G A701V | 10.94% |
| B.1.617.1 | Kappa | L452R E484Q D614G P681R | 18.34% |
| C.37 | Lambda | G75V T76I R246- S247- Y248- L249- T250- P251- G252- D253N L452Q F490S D614G T859N | 34.22% |
| C.37* | Lambda | G75V T76I L452Q F490S D614G T859N | 16.88% |
| BA.1 | Omicron | A67V H69- V70- T95I G142D V143- Y144- Y145- N211I L212V ins214EPE G339D S371L S373P S375F K417N N440K G446S S477N T478K E484A Q493R G496S Q498R N501Y Y505H T547K D614G H655Y N679K P681H N764K D796Y N856K Q954H N969K L981F | 97.52% |
| BA.2 | Omicron | T19I L24- P25- P26- A27S G142D V213G G339D S371L S373P S375F T376A D405N R408S K417N N440K S477N T478K E484A Q493R Q498R N501Y Y505H D614G H655Y N679K P681H N764K D796Y Q954H N969K | 91.16% |
| BA.4/5 | Omicron | T19I L24- P25- P26- A27S H69- V70- G142D V213G G339D S371F S373P S375F T376A D405N R408S K417N N440K L452R S477N T478K E484A F486V Q498R N501Y Y505H D614G H655Y N679K P681H N764K D796Y Q954H N969K | 94.38% |
| A.VOI.V2 | | D80Y Y144- I210- D215G R246- S247- Y248- L249M W258L R346K T478R E484K H655Y P681H Q957H | 64.54% |
| B.1.1.298 | | Y453F D614G I692V M1229I | 5.42% |
| B.1.160 | | S477N S494P D614G K1191N | 3.81% |
| B.1.258 | | H69- V70- L189F N439K D614G V772I | -9.28% |
| B.1.517 | | G181V G252V N501T D614G P812L | -7.87% |

**Table S7. Correlation between Epitope Alteration Scores and $pVNT_{50}$ Reduction per Antibody Class.** We report the correlation between epitope alteration scores using all available epitopes until October 1st 2021 and $pVNT_{50}$ reduction when considering different classes of antibodies, defined in Barnes *et al.* [8]. For instance, when limiting to class 1, the epitope alteration scores are calculated using only antibodies of class 1. Among the 227 antibodies in collected data, 88 have formally defined antibody classes. Despite using fewer antibodies, a strong correlation is observed and limiting antibody classes often leads to poorer correlation.

| Antibody Class | Pearson *r* | Pearson *p*-value | Spearman *r* | Spearman *p*-value |
|---|---|---|---|---|
| All four classes | 0.63 | 2E-03 | 0.74 | 7E-05 |
| Class 1 | 0.49 | 2E-02 | 0.68 | 5E-04 |
| Class 2 | 0.69 | 4E-04 | 0.74 | 7E-05 |
| Class 3 | 0.62 | 2E-03 | 0.59 | 4E-03 |
| Class 4 | 0.61 | 3E-03 | 0.63 | 2E-03 |
| Exclude Class 1 | 0.72 | 2E-04 | 0.79 | 1E-05 |
| Exclude Class 2 | 0.59 | 4E-03 | 0.68 | 5E-04 |
| Exclude Class 3 | 0.56 | 7E-03 | 0.71 | 2E-04 |
| Exclude Class 4 | 0.64 | 1E-03 | 0.71 | 2E-04 |

**Table S8. Correlation between Epitope Alteration Scores and $pVNT_{50}$ Reductions per Epitope Class.** We report the correlation between epitope alteration scores using all available epitopes until October 1st 2021 and $pVNT_{50}$ reduction when considering different classes of epitopes defined in Sikora et al.[7]. For instance, when limiting to class E1, the epitope alteration scores are calculated based only on mutations in the positions of class E1. For certain cases, when limiting epitopes, all sequences share the same epitope alteration score and therefore the correlation is not available.

| Epitope Class | Pearson r | Pearson p-value | Spearman r | Spearman p-value |
|---|---|---|---|---|
| All | 0.64 | 1E-03 | 0.77 | 3E-05 |
| E1 | 0.26 | 3E-01 | 0.16 | 5E-01 |
| E2 | 0.60 | 3E-03 | 0.54 | 9E-03 |
| E3 | 0.39 | 7E-02 | 0.59 | 4E-03 |
| E4 | 0.44 | 4E-02 | 0.60 | 3E-03 |
| E5 | 0.59 | 4E-03 | 0.61 | 2E-03 |
| E6 | 0.61 | 2E-03 | 0.59 | 4E-03 |
| E7 | N/A | N/A | N/A | N/A |
| E8 | N/A | N/A | N/A | N/A |
| E9 | N/A | N/A | N/A | N/A |
| Exclude E1 | 0.67 | 6E-04 | 0.79 | 1E-05 |
| Exclude E2 | 0.66 | 9E-04 | 0.78 | 2E-05 |
| Exclude E3 | 0.58 | 5E-03 | 0.73 | 1E-04 |
| Exclude E4 | 0.63 | 2E-03 | 0.76 | 4E-05 |
| Exclude E5 | 0.67 | 7E-04 | 0.76 | 4E-05 |
| Exclude E6 | 0.71 | 2E-04 | 0.76 | 4E-05 |
| Exclude E7 | 0.64 | 1E-03 | 0.77 | 3E-05 |
| Exclude E8 | 0.64 | 1E-03 | 0.77 | 3E-05 |
| Exclude E9 | 0.64 | 1E-03 | 0.77 | 3E-05 |

**Table S9. Correlation between Epitope Alteration Scores and $pVNT_{50}$ Reductions per Epitope Region.** We report the correlation between epitope alteration scores using all available epitopes until October 1st 2021 and $pVNT_{50}$ reduction when considering different regions of epitopes. For instance, when limiting to the RBD region, the epitope alteration scores are calculated based only on mutations in the RBD region. For certain regions, when limiting epitopes, all sequences share the same epitope alteration score and therefore the correlation is not available. For all the sequences with $pVNT_{50}$ data, the mutations are defined on S1 and the best correlation is observed using all regions including NTD and RBD.

| Epitope Region | Pearson r | Pearson p-value | Spearman r | Spearman p-value |
|---|---|---|---|---|
| All | 0.64 | 1E-03 | 0.77 | 3E-05 |
| NTD | 0.54 | 9E-03 | 0.59 | 4E-03 |
| RBD | 0.62 | 2E-03 | 0.73 | 1E-04 |
| S1 | 0.64 | 1E-03 | 0.77 | 3E-05 |
| S2 | N/A | N/A | N/A | N/A |
| Exclude NTD | 0.62 | 2E-03 | 0.73 | 1E-04 |
| Exclude RBD | 0.54 | 9E-03 | 0.59 | 4E-03 |
| Exclude S1 | N/A | N/A | N/A | N/A |
| Exclude S2 | 0.64 | 1E-03 | 0.77 | 3E-05 |

**Table S10. Early detection of variants of concerns.** The summary table shows that, with 12 sequences per week, the EWS can detect WHO designated variants months before the WHO official designation date. The average lead time for early detection across is 52 days. The average number of submissions is 9697 and 66 for WHO designation and EWS detection correspondingly.

| **WHO Label** | **Lineage** | **WHO Designation Date** | **EWS Detection Date** | **First Submission Date** | **Lead Time (Days)** | **Number of submissions upon WHO Designation** | **Number of submissions upon EWS Detection** |
|---|---|---|---|---|---|---|---|
| Alpha | B.1.1.7 | 2020-12-18 | 2020-11-25 | 2020-11-25 | 23 | 54 | 1 |
| Beta | B.1.351 | 2020-12-18 | 2020-11-26 | 2020-11-26 | 22 | 207 | 2 |
| Gamma | P.1 | 2021-01-11 | 2021-01-10 | 2021-01-10 | 1 | 4 | 4 |
| Delta | B.1.617.2 | 2021-05-11 | 2021-04-28 | 2021-03-08 | 13 | 2774 | 492 |
| Epsilon | B.1.429 | 2021-03-05 | 2020-11-21 | 2020-11-21 | 104 | 10386 | 1 |
| Zeta | P.2 | 2021-03-17 | 2020-12-20 | 2020-12-02 | 87 | 1360 | 41 |
| Eta | B.1.525 | 2021-03-17 | 2021-01-04 | 2021-01-04 | 72 | 652 | 2 |
| Theta | P.3 | 2021-03-24 | 2021-02-25 | 2021-02-25 | 27 | 94 | 1 |
| Iota | B.1.526 | 2021-03-24 | 2021-01-11 | 2020-12-09 | 72 | 3544 | 10 |
| Kappa | B.1.617.1 | 2021-04-04 | 2021-03-31 | 2021-03-05 | 4 | 123 | 116 |
| Lambda | C.37 | 2021-06-14 | 2021-02-26 | 2021-02-15 | 108 | 1402 | 4 |
| Mu | B.1.621 | 2021-08-30 | 2021-03-29 | 2021-03-11 | 154 | 4475 | 2 |
| Omicron | BA.1 | 2021-11-26 | 2021-11-25 | 2021-11-22 | 1 | 95 | 73 |
| Omicron | BA.2 | 2022-02-22 | 2021-11-27 | 2021-11-27 | 87 | 127546 | 1 |
| Omicron | BA.4 | 2022-05-18 | 2022-04-13 | 2022-03-08 | 35 | 1549 | 123 |
| Omicron | BA.5 | 2022-05-18 | 2022-04-26 | 2022-03-15 | 22 | 883 | 181 |

**Table S11**. Comparison between EWS detection capabilities and three baselines. Two baselines are based on unsupervised learning (UMAP) and one baseline is supervised (GLM). N/A means the method failed to detect the variant. Negative values indicate detection after WHO designation.

| HRV | UMAP | GLM | Growth Score | Epitope Alteration Score | Semantic Change Score | Immune Escape Score |
|---|---|---|---|---|---|---|
| Alpha | 23 | -70 | 23 | 23 | 23 | 23 |
| Beta | 21 | 11 | 21 | 22 | 22 | 22 |
| Gamma | -101 | 1 | -1 | 1 | -120 | 1 |
| Delta | 2 | -2 | 36 | -74 | 34 | 5 |
| Epsilon | 22 | 23 | 104 | 59 | 104 | 104 |
| Zeta | -21 | 70 | 87 | 105 | -14 | 87 |
| Eta | 41 | 55 | 42 | -27 | 33 | 72 |
| Theta | -68 | N/A | 21 | 27 | 27 | 27 |
| Iota | -42 | 79 | 15 | 72 | 0 | 72 |
| Kappa | -19 | -305 | 20 | N/A | 4 | 4 |
| Lambda | -108 | N/A | 70 | 60 | 108 | 108 |
| Mu | 71 | N/A | 140 | 154 | 131 | 154 |
| Omicron (BA.1) | -7 | -40 | 1 | 4 | 1 | 1 |
| Omicron (BA.2) | 28 | -9 | 87 | 87 | 42 | 87 |
| Omicron (BA.4) | 30 | 34 | 35 | 30 | 35 | 35 |
| Omicron (BA.5) | 7 | 26 | 35 | 15 | 15 | 22 |
| Average | -8 | -10 | 46 | 37 | 28 | 52 |

#### Supplementary Information References
